# A novel family of sugar-specific phosphodiesterases that remove zwitterionic modifications of *N*-acetylglucosamine

**DOI:** 10.1101/2023.06.01.543256

**Authors:** Samantha L. Fossa, Brian P. Anton, Daniel W. Kneller, Laudine M.C. Petralia, Madison L. Boisvert, Mehul B. Ganatra, Saulius Vainauskas, S. Hong Chan, Cornelis H. Hokke, Jeremy M. Foster, Christopher H. Taron

**Affiliations:** New England Biolabs, 240 County Road, Ipswich, MA 01938, USA; Department of Parasitology, Leiden University – Center of Infectious Diseases, Leiden University Medical Center, Leiden, The Netherlands

**Keywords:** Phosphorylcholine, phosphodiesterase, functional metagenomics, parasitology, glycobiology, glycan analysis, N-glycans, *N*-acetylglucosamine (GlcNAc), glycosphingolipid

## Abstract

The zwitterions phosphorylcholine (PC) and phosphoethanolamine (PE) are often found esterified to certain sugars in polysaccharides and glycoconjugates in a wide range of biological species. One such modification involves PC attachment to the 6-carbon of *N*-acetylglucosamine (GlcNAc-6-PC) in N-glycans and glycosphingolipids (GSLs) of parasitic nematodes, a modification that helps the parasite evade host immunity. Knowledge of enzymes involved in the synthesis and degradation of PC and PE modifications is limited. More detailed studies on such enzymes would contribute to a better understanding of the function of PC modifications and have potential application in the structural analysis of zwitterion-modified glycans. In this study, we used functional metagenomic screening to identify phosphodiesterases encoded in a human fecal DNA fosmid library that remove PC from GlcNAc-6-PC. A novel bacterial phosphodiesterase was identified and biochemically characterized. This enzyme (termed GlcNAc-PDase) shows remarkable substrate preference for GlcNAc-6-PC and GlcNAc-6-PE, with little or no activity on other zwitterion-modified hexoses. The identified GlcNAc-PDase protein sequence is a member of the large endonuclease/exonuclease/phosphatase (EEP) superfamily where it defines a distinct subfamily of related sequences of previously unknown function, mostly from *Clostridium* bacteria species. Finally, we demonstrate use of GlcNAc-PDase to confirm the presence of GlcNAc-6-PC in N-glycans and GSLs of the parasitic nematode *Brugia malayi* in a glycoanalytical workflow.

## INTRODUCTION

The outer surface of cells is coated with a diverse set of sugar-based polymers called glycans. Glycans are assembled from monosaccharide building blocks into long polysaccharide chains or into smaller but structurally complex oligosaccharides covalently attached to proteins or lipids (glycoconjugates). Adding to this complexity, site-specific enzymatic modifications of glycans with different chemical groups (*e.g*., methyl, sulfate, phosphate, acetyl, or zwitterions) may occur. These modifications are often referred to as “post-glycosylational modifications” (PGMs) (1, 2). PGMs act in many contexts as important modulators of glycan biological function. Changes in the PGM content of cellular or tissue glycomes have been observed in many diseases including autoimmune disorders, cystic fibrosis, infections, and certain cancers (2). As such, this area of glycobiology has enormous potential for therapeutic and molecular diagnostic application. However, analysis of PGMs remains technically challenging, in part due to there being few well-characterized and highly selective molecular tools (*e.g*., enzymes, binding proteins) that assist in their detection and simplify their analysis.

One interesting class of PGM consists of zwitterionic modification of glycans. These PGMs involve esterification of phosphorylcholine (PC) or phosphoethanolamine (PE) to a sugar ring (Fig. 1A). They are present on a variety of polysaccharides and glycoconjugates throughout biology. In eukaryotes, PC and PE have been observed on N-glycans from several invertebrates (3) and *Penicillium* fungi (4). They are prevalent on N-glycans and glycolipids of nematodes, including several mammalian parasite species (5–10), where they help the parasite evade host immunity through inhibition of immune cell activity (6). Additionally, the eukaryotic glycosylphosphatidylinositol (GPI) anchor possesses a PE moiety that participates in linking the GPI to protein, and further side-branching PE residues decorate its glycan core in mammals and yeasts (11–14). In bacteria, PE has been found on the cellulosic biofilm of *Escherichia coli* where it likely protects the polymer from degradation by hydrolytic enzymes (15). Additionally, sugar-associated zwitterions have been observed on diacylglycerol in *Clostridium tetani* (16), and in cell wall polysaccharides and teichoic acids of several pathogenic bacteria (17, 18). Despite the breadth of PC/PE distribution in glycobiology, few enzymes that either add (12, 19, 20) or remove (21–23) these groups from sugars have been defined. Thus, better definition of the range of enzymes involved in manipulating glycan zwitterions will lead to better understanding of the biosynthesis, degradation, and functions of these PGMs, and may also have utility in glycoanalytical workflows. The primary aim of the current study was to use high-throughput functional metagenomic screening to discover enzymes that hydrolyze zwitterions from sugars. Functional metagenomics is a powerful method for detecting enzymatic activities encoded in environmental DNA (eDNA). A typical approach involves eDNA isolation from any environment, cloning large eDNA fragments into fosmid vectors (typically in *Escherichia coli*), and screening clones in high throughput for microexpression of a desired enzyme activity. The main advantages of this method are that enzymes are identified based on their activity (not similarity to known protein families) and that proteins from both cultivable and uncultivable microorganisms can be found. Thus, functional metagenomics is well-suited to identify new specificities and often new protein families.

We previously established a functional metagenomic screen to identify enzymes that hydrolyze PGMs, specifically the removal of sulfate from C6 of *N*-acetylglucosamine (GlcNAc) (24). In the current study, we have further adapted this general strategy to seek enzymes that hydrolyze PC from C6 of GlcNAc (GlcNAc-6-PC). This structure is the predominant zwitterion modification of N-glycans and glycolipids in the human parasitic nematode *Brugia malayi* (9) and other filarial parasites. We report successful identification and biochemical characterization of a new family of phosphodiesterases with remarkable selectivity for zwitterion modifications of GlcNAc at C6. We define the biochemical properties and preliminary structural features of multiple members of this protein family. Finally, we show the utility of this enzyme for detection of GlcNAc-6-PC in *B. malayi* N-glycans and glycolipids. Our study establishes the feasibility of using functional metagenomics to identify enzymes that act on zwitterion-modified sugars, defines a new family of sugar-specific phosphodiesterases, and validates an important new enzyme specificity for use in glycoanalytical workflows.

**Figure 1.**
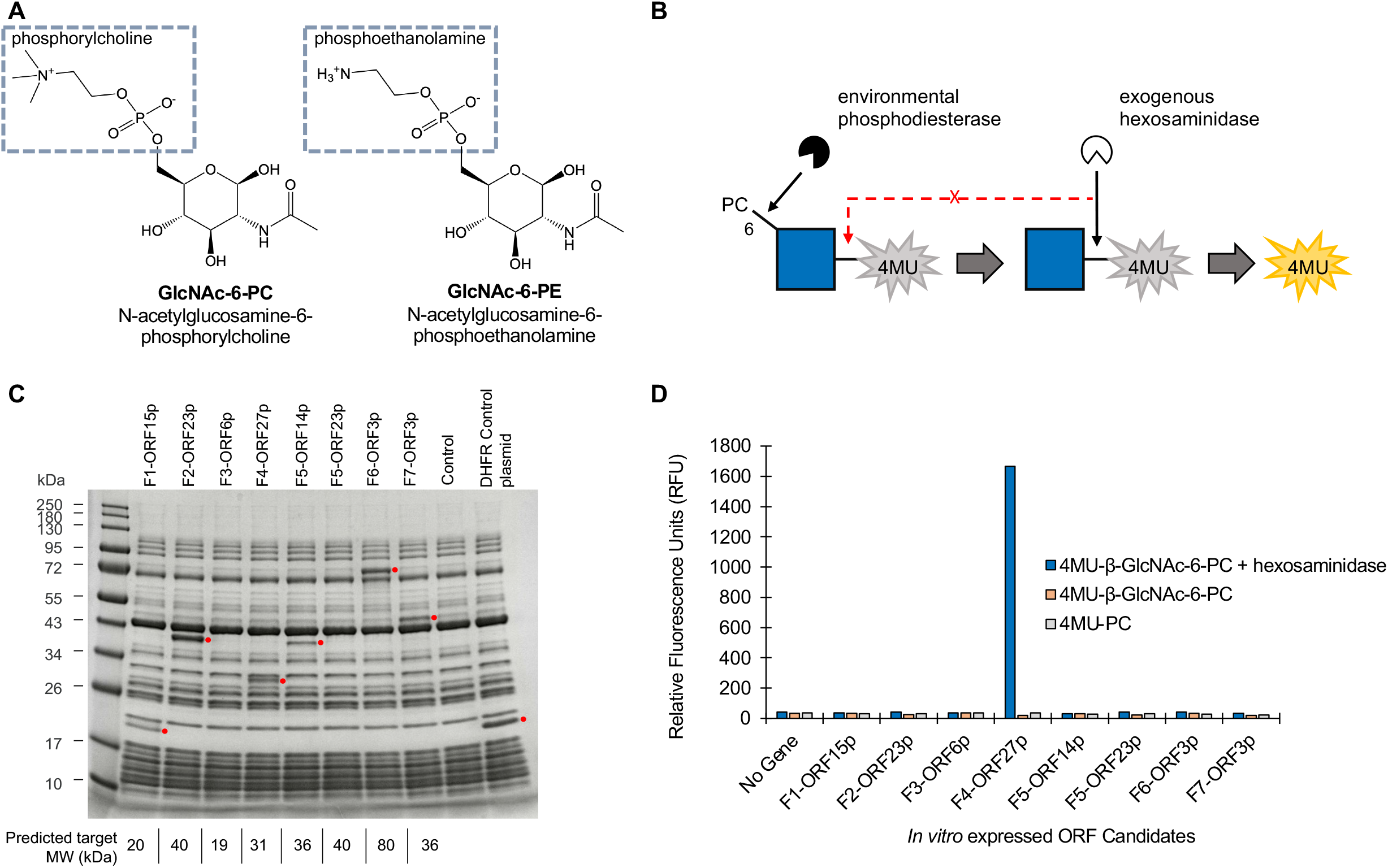
Metagenomic screening strategy and candidate phosphodiesterase assessment. (A) Chemical structures of GlcNAc-6-PC and GlcNAc-6-PE. (B) A schematic of the coupled assay used to screen for environmental GlcNAc-6-PC phosphodiesterases. A quenched fluorescent GlcNAc analog (4MU-β-GlcNAc-6-PC) is used as substrate. A fluorescent signal (yellow 4-MU) is generated upon sequential removal of PC by an environmental phosphodiesterase and GlcNAc by β-*N*-Acetylhexosaminidasef provided in the assay. Screening data is provided in Figure S1. Glycan structure is represented using the SNFG nomenclature. (C) SDS-PAGE separation of *in vitro* expressed proteins from 8 candidate ORFs identified in the DNA sequence of fosmid “hits” (see Table 1). Shown are expressed proteins (red circles) and the predicted molecular weight (kDa) of each (bottom). (D) Expressed proteins were assayed for phosphodiesterase activity using 4MU-β-GlcNAc-6-PC (+/-hexosaminidase) and 4MU-PC substrates. Fluorescence was measured at λex=365 nm and λem=445 nm after 3 h.

## RESULTS

### A functional metagenomic screen for GlcNAc-6-PC phosphodiesterases

A functional metagenomic screen was designed to find enzymes able to remove PC from C6 of GlcNAc. The assay used for screening was adapted from one we previously used to identify sulfatases that remove sulfate from C6 of GlcNAc (24). In the present version, we used a fluorogenic substrate consisting of GlcNAc with PC esterified to C6 and 4-methylumbelliferone (4-MU) attached to C1 (4MU-ý-GlcNAc-6-PC). This assay permits an expressed environmental phosphodiesterase to liberate PC from 4MU-ý-GlcNAc-6-PC, after which an exogenous ý-hexosaminidase provided in the assay mixture hydrolyzes 4MU-ý-GlcNAc to generate a fluorescent signal (Fig. 1B). A previously reported library of fosmid clones containing ∼40 kb inserts of human fecal eDNA was used for screening (24).

A total of 6,144 fosmid clones were screened resulting in identification of 23 “hits”. Hits were defined as clones yielding an assay signal at least 10 standard deviations above the mean background fluorescence measured for clones carrying only the empty pCC1Fos fosmid vector (Fig. S1). Fosmid DNA was isolated from each hit and subjected to long-read nucleotide sequencing as described in Materials and Methods. The assembled nucleotide sequence of 7 fosmid inserts, termed F1-F7, were deposited in GenBank under the accession numbers OQ439824-OQ439830. Each fosmid sequence was analyzed by computational prediction of encoded open reading frames (ORFs) using MetaGeneMark and BLASTP analysis of each ORF. In our prior screen for enzymes that remove sulfate from GlcNAc-6-SO4, we identified both GlcNAc-specific sulfatases and a hexosaminidase that preferentially hydrolyzes GlcNAc-6-SO4 (24). As such, in this study, we focused on 9 fosmid ORFs having annotations that suggested possible roles in either manipulation of phosphate or sugar hydrolysis (Table 1). Of these 9 ORFs, 8 were successfully amplified and subsequently expressed *in vitro* using PURExpress^®^ (Fig. 1C), and the reactions were assayed on 4MU-ý-GlcNAc-6-PC, both with and without exogenous hexosaminidase, and 4MU-PC (Fig. 1D). A single protein encoded by ORF27 on fosmid F4 was clearly active on 4MU-ý-GlcNAc-6-PC in the presence of exogenous hexosaminidase but showed no activity in its absence or on 4MU-PC (herein referred to as GlcNAc-PDase for this study). In an orthologous analysis, the enzyme was shown by ultra-performance liquid chromatography with inline fluorescence and mass detection (UPLC-FLR-MS) to liberate PC from 4MU-ý-GlcNAc-6-PC (Fig. S2), confirming that it was functioning as a phosphodiesterase. Characterization of the biochemical properties of GlcNAc-PDase and its protein family is the subject of the remainder of this study.

### Analysis of the GlcNAc-PDase protein sequence and sequence family

Basic features of the deduced GlcNAc-PDase sequence were analyzed computationally. The transmembrane prediction algorithm TMHMM 2.0 and the signal peptide prediction tool SignalP 5.0 revealed that GlcNAc-PDase lacks transmembrane segments and possesses a signal peptide (predicted cleavage after glycine-34: VPGτSG), suggesting it is likely a soluble secreted protein in its native environment. Protein BLAST against the GenBank non-redundant database revealed that the two most similar proteins were both from Clostridiales bacteria (accession numbers MBS6702114 and MBS6250406, with 99% and 94% identity, respectively). These uncharacterized proteins are members of a very large endonuclease/exonuclease/phosphatase (EEP) superfamily (Pfam protein family PF03372 and Conserved Domain Database family cd08372). Characterized proteins in this diverse enzyme superfamily share a common catalytic mechanism that breaks phosphodiester bonds. Members include enzymes that act upon nucleic acids such as certain AP endonucleases and DNase I exonucleases, and enzymes that act upon certain phosphodiester-linked lipids like sphingomyelin.

A sequence clustering analysis was performed to determine the relatedness of GlcNAc-PDase to 900 other sequences from the EEP superfamily. GlcNAc-PDase clusters closely with groupings of protein sequences (cd09079, cd09083, cd09084) of unknown function and sphingomyelinases (cd09078) (Fig. 2A). Within the immediate GlcNAc-PDase cluster, only two proteins, yeast Cwh43p and human PGAP2IP, have been previously studied (25, 26). These two related proteins have been implicated in lipid remodeling events during biosynthesis of glycosylphosphatidylinositol (GPI) anchors, however, the precise role of their EEP domains is still vague. Thus, sequence cluster analysis points to the GlcNAc-PDase group as being a unique and distinct subfamily of PF03372.

**Figure 2.**
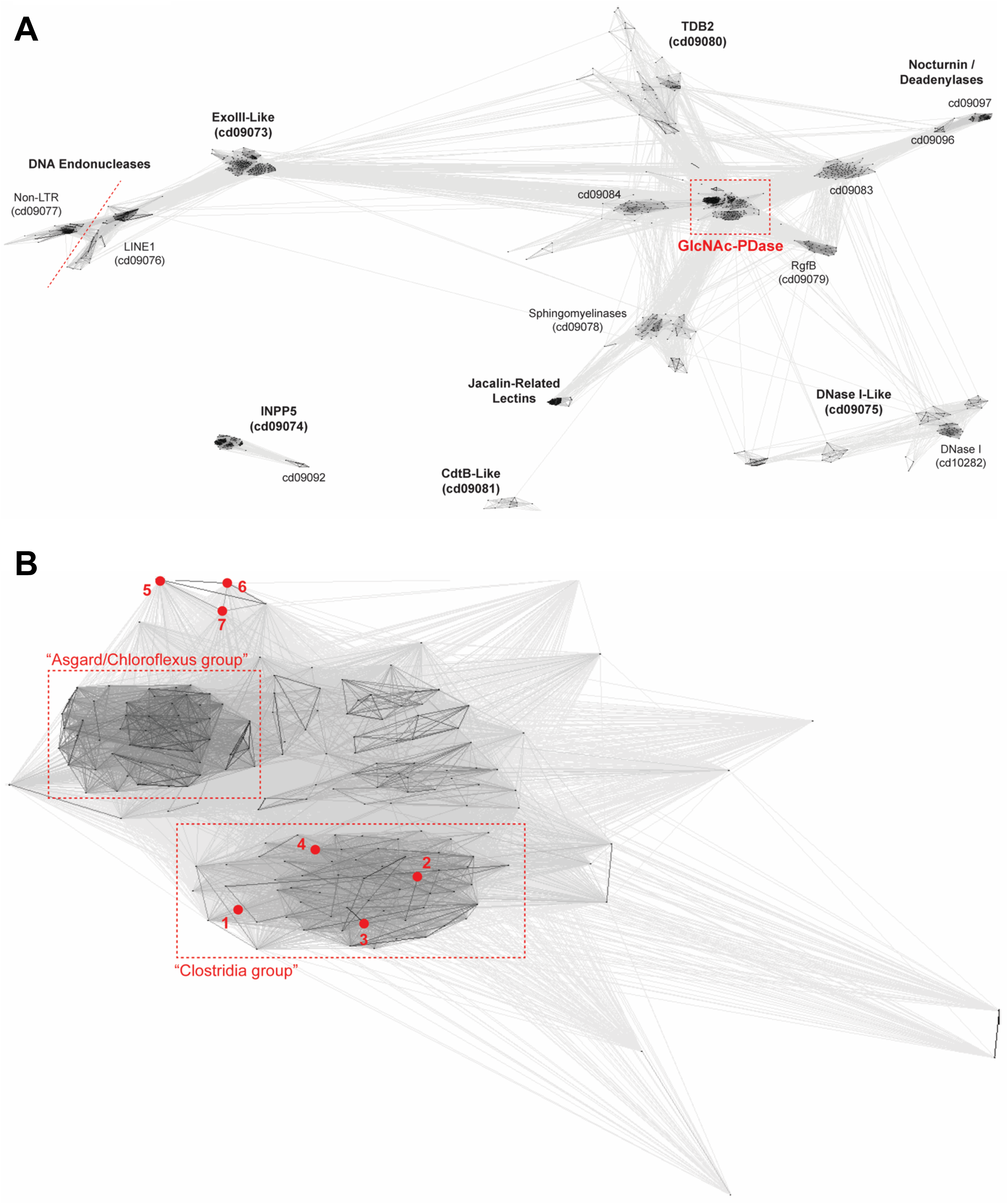
GlcNAc-PDase sequence clustering analysis within EEP superfamily. Results of sequence clustering of EEP protein sequences using CLANS (38). Each protein is represented by a node (point). Edges are drawn between nodes with sufficient pairwise sequence similarity, as determined by BLAST *P*-value; in general, the darker the edge, the greater the similarity. Groups of proteins with a mutually high degree of relatedness will tend to cluster. (A) Clustering of 900 proteins from throughout the EEP superfamily. Clusters are labeled with the family name based on characterized and/or named members and, where defined, the CDD family number. Full-length protein sequences are used, and so for multidomain proteins, some edges may be formed or strengthened based on similarity outside the EEP domain. The red box indicates the GlcNAc-PDase group, a unique and defined cluster that does not have an assigned CDD number. (B) Clustering of 186 proteins from the GlcNAc-PDase group, now trimmed to restrict the sequences to the EEP domain only. Two taxonomically specific clusters are indicated by the red boxes, one comprising sequences primarily from Asgardarchaeota (archaea) and Chloroflexota (bacteria), and the other comprising sequences primarily from Clostridia (bacteria). Seven nodes (proteins) are labeled by number, including the four proteins experimentally characterized in this work [#1-4] and three previously studied eukaryotic proteins [#5-7]: [1] GlcNAc-PDase from an unknown strain, [2] HCG67795 from Clostridiales bacterium, [3] MBE6669927 from Ruminococcaceae bacterium, [4] NLD61619 from Candidatus Sumerlaeota bacterium, [5] Cwh43p from *Saccharomyces cerevisiae*, [6] Cwh43p from *Schizosaccharomyces pombe*, and [7] PGAP2IP from *Mus musculus*.

The GlcNAc-PDase group was further stratified by clustering its 187 sequences (Fig. 2B). Two prominent, taxonomically restricted subclusters, one including mostly proteins from Asgardarchaeota (archaea) and Chloroflexota (bacteria), and the other including mostly proteins from Clostridia (bacteria), are noted. However, the functional significance, if any, of the subclusters within the GlcNAc-PDase group is unclear. The “Clostridia group”, including those with the highest similarity to GlcNAc-PDase, comprises 65 sequences. Most member proteins are 219-357 amino acids long and are composed nearly entirely of a single EEP domain with a secretion peptide and no transmembrane segments. While not biochemically explored in this study, proteins in the “Asgard/Chloroflexus group” were divergent from the Clostridia group in their size and domain organization.

### Biochemical properties of GlcNAc-PDase

To define the basic biochemical properties of GlcNAc-PDase, its recombinant production and purification was undertaken. His-tagged GlcNAc-PDase was expressed in *E. coli* and purified using two rounds of nickel affinity chromatography as described in Experimental Procedures (Fig. S3). Purified GlcNAc-PDase required a divalent metal ion for catalysis *in vitro* (Fig. 3A). It was most active in the presence of Mg^2+^ and Mn^2+^ ions, and to a lesser degree, Ni^2+^ and Co^2+^ ions. The enzyme was active over a broad temperature range (15-50 °C) with an optimum at 37 °C (Fig. 3B) and was active from pH 6-10 (Fig. 3C). Applying these biochemical properties, a unit was defined as the amount of enzyme needed to liberate PC from 10 nmol of GlcNAc-6-PC in 50 mM Tris-HCl, pH 8.0 containing 5 mM MgCl2 in 1 h at 37 °C.

**Figure 3.**
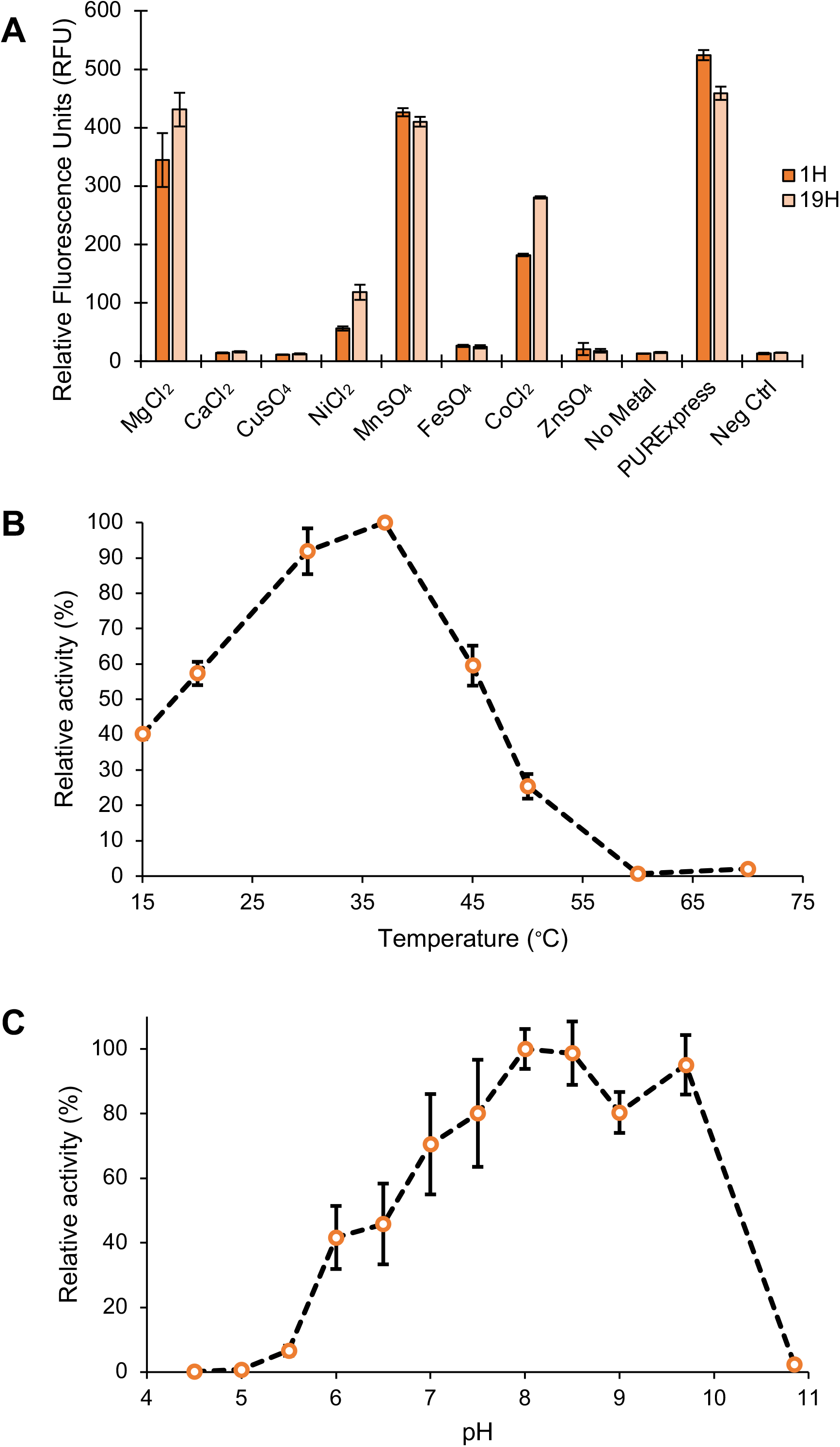
GlcNAc-PDase biochemical characterization. *In vivo* purified GlcNAc-PDase was used to define the enzyme’s catalytic properties. (A) The substrate 4MU-ý-GlcNAc-6-PC was used to determine GlcNAc-PDase activity in the presence of metal ions. Procainamide labeled GlcNAc-6-PC was used as substrate to determine GlcNAc-PDase optimal (B) temperature and (C) pH. All reactions were performed in triplicate. PURExpress^®^: GlcNAc-PDase expressed *in vitro*; Neg Ctrl: PURExpress^®^ mixture without DNA template.

### Specificity of selected “Clostridia group” proteins

The specificity of GlcNAc-PDase and other selected proteins from the Clostridia group was explored *in vitro*. For this experiment, three proteins from the Clostridia group were selected for functional comparison to GlcNAc-PDase. These proteins are referred to by their GenBank accession numbers as follows: NLD61619, from Candidatus Sumerlaeota bacterium; MBE6669927, from *Oscillospiraceae* bacterium; and HCG67795, from Clostridiales bacterium.

These proteins consist of 262-297 amino acids, have a single EEP domain, and share 34%, 38%, and 38% identity with GlcNAc-PDase, respectively (See Supporting Information Fig. S4 for additional information). Expression of each was achieved by *in vitro* transcription and translation (IVTT) using the PURExpress^®^ system (Fig. S5A). IVTT-produced material was assayed for activity on the screening substrate 4MU-ý-GlcNAc-6-PC with ý-hexosaminidase to confirm that it was functional (Fig. S5B). The specificity of the four enzymes was explored using a panel of synthetic monosaccharides possessing zwitterions. In these assays, IVTT-produced enzyme was incubated with a monosaccharide substrate, after which, the sugar was reductively labeled with procainamide. The reaction products were then analyzed by UPLC-FLR. The ability of each enzyme to remove PC or PE from 8 different substrates was determined (Table 2; Fig. S6-S13). The four enzymes performed similarly with a marked preference for removal of PC or PE from the 6-carbon of GlcNAc (Fig. 4A, 4B, S6, and S10). Significantly lower hydrolysis of Glc-6-PC was observed for all four proteins, and only related proteins 1 (NLD61619) and 2 (MBE6669927) showed lower hydrolysis of Glc-6-PE (Fig. 4C, 4D, S7, and S11). All four enzymes were each unable to efficiently remove PC or PE from the 6-carbon of galactose or mannose, or PE from the 2-carbon of mannose (a linkage found in fungal and mammalian GPI anchors) (Fig. S8, S9, S12, and S13). These data support the conclusion that the Clostridia group cluster of proteins represents a family of highly specific GlcNAc-preferring phosphodiesterases.

**Figure 4.**
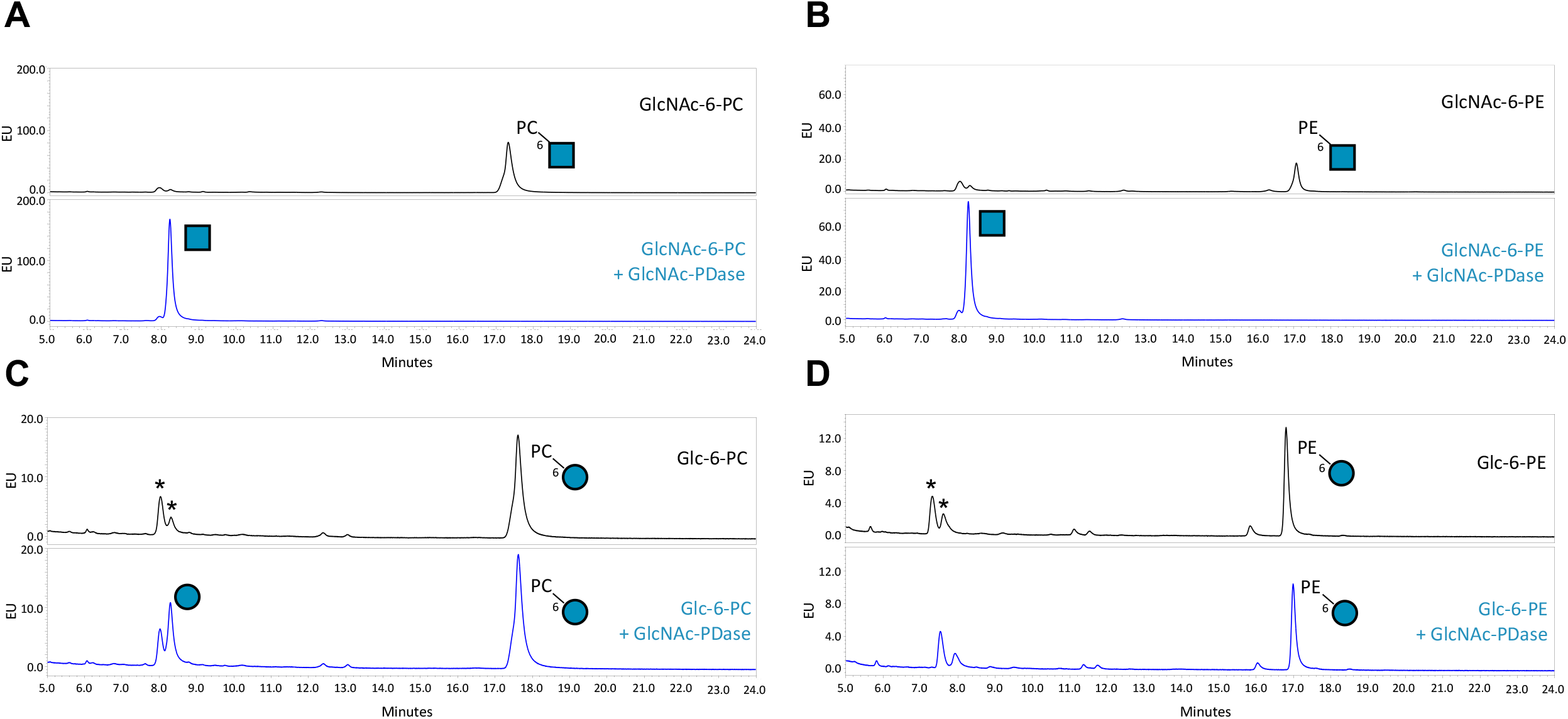
GlcNAc-PDase specificity analyzed by UPLC-FLR. *In vitro* expressed GlcNAc-PDase was incubated with (A) GlcNAc-6-PC, (B) GlcNAc-6-PE, (C) Glc-6-PC, or (D) Glc-6-PE for 1 h, procainamide labeled, then separated by UPLC-FLR. For each set of traces, the top panel shows a control reaction with substrate alone and the bottom panel shows digestion of the substrate by GlcNAc-PDase. Glycan structures are represented using the SNFG nomenclature: blue circle = glucose, blue square = GlcNAc, PC = phosphorylcholine and PE = phosphoethanolamine. Numbers reflect the sugar carbon that is bonded with the zwitterion. Peaks detected due to the intrinsic fluorescence of the PURExpress^®^ components are denoted with an asterisk (*)

### Structural modeling of GlcNAc-PDase

The tertiary structure of GlcNAc-PDase was predicted with AlphaFold2 to provide insight into the catalytic structure-function relationship (Fig. 5A). The top 5 models displayed consistent and confident scores in the Predicted Local Distance Difference Test (pLDDT) (27) suggesting reliable backbone predictions for most of the protein and permitting investigation of active site residues.

**Figure 5.**
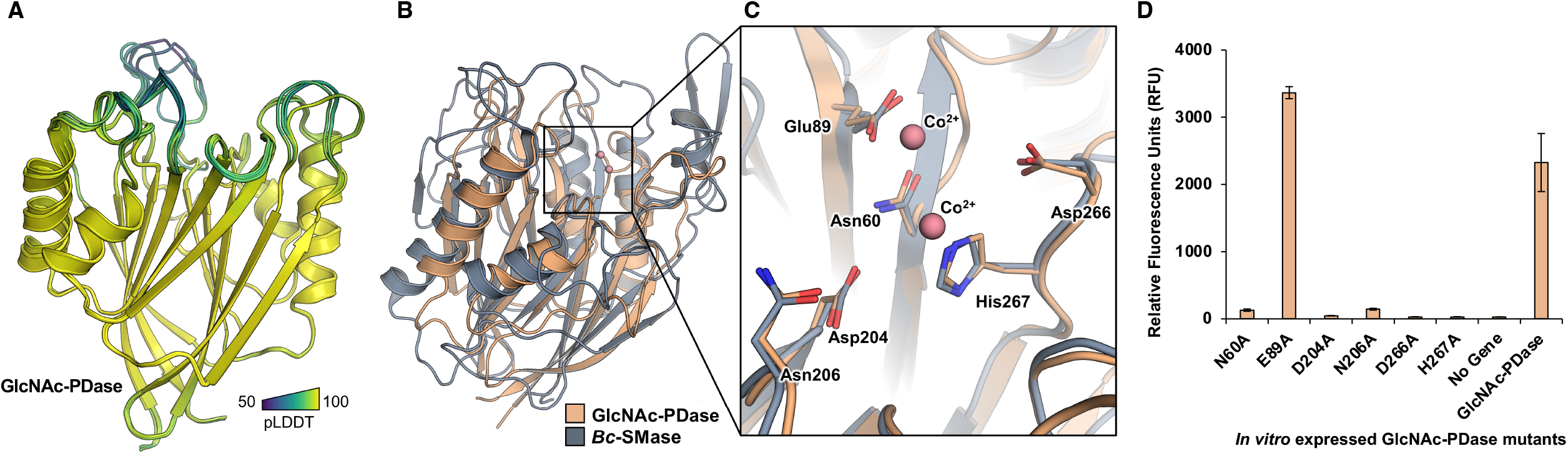
Structural analysis of GlcNAc-PDase. (A) Predicted structure of the GlcNAc-PDase sequence in cartoon representation colored in Viridis by Predicted Local Distance Difference Test (pLDDT) scores. The structured regions from residues 50-278 for ranks 1-5 are shown. For rank 1, per-residue estimate confidence scores for the structured regions from residues 50-278 have a mean ± standard deviation of 96.0 ± 4.4. (B) Superposition of the GlcNAc-PDase predicted structure (light orange) with a 1.8Å X-ray crystal structure of *B. cereus* sphingomyelin phosphodiesterase (*Bc-*SMase) in complex with two cobalt ions (blue-grey, Co^2+^ as salmon spheres). (C) Metal ion binding site of GlcNAc-PDase and *Bc-* SMase. Residues are shown as sticks with the GlcNAc-PDase sequence used for position numbers. (D) Activity of GlcNAc-PDase metal binding site alanine mutants. Six mutants of GlcNAc-PDase (N60A, E89A, D04A, N206A, D266A, and H267A) were expressed *in vitro*. For each mutant, fluorescence release from 4MU-ý-GlcNAc-6-PC plus an exogenous hexosaminidase was assessed after 1 h in triplicate reactions. Mutants were compared to a no gene control and wild-type GlcNAc-PDase.

**Figure 6.**
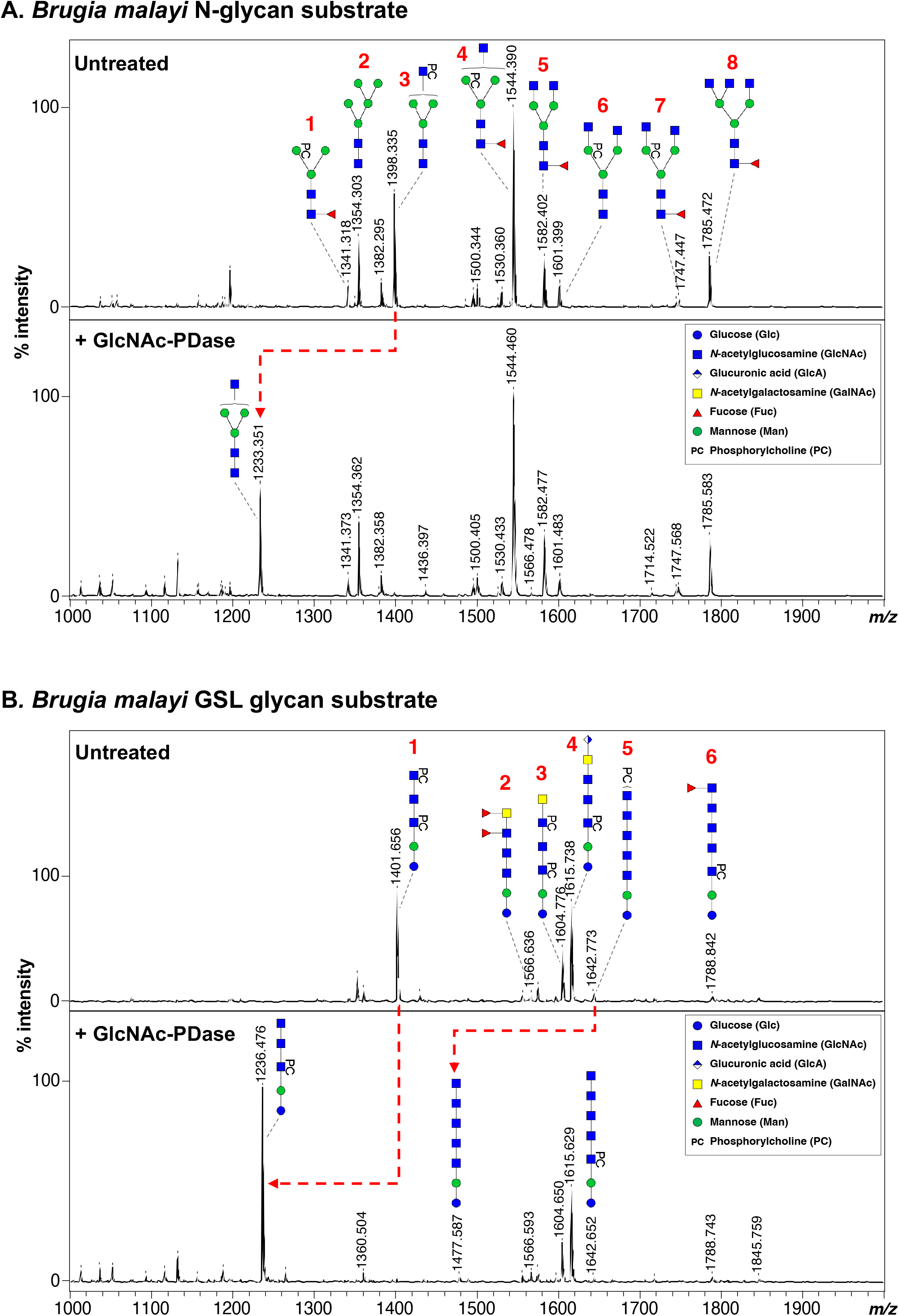
Digestion of *B. malayi* glycans with GlcNAc-PDase on MALDI-TOF-MS. Enzymatically released, AA-labeled and UHPLC purified *B. malayi* (A) N-linked and (B) GSL glycans were subjected to digestion with GlcNAc-PDase. Resulting digestion products are highlighted using red dashed arrows. MALDI-TOF-MS spectra monoisotopic masses are indicated for glycan ions and glycan structures are represented using the CFG nomenclature. See Symbol key inset. MALDI-TOF-MS raw data can be found in Table S2.A.

The predicted coordinates for GlcNAc-PDase were first queried against experimental structures to identify homologous folds using the DALI protein structure comparison server (28, 29). Over 50 X-ray structures in the PDB90 representative database returned high Z-scores (>10) and low sequence identity (<25%). An X-ray crystal structure of Sphingomyelin phosphodiesterase from *Bacillus cereus* (*Bc*-SMase, PDB 2DDS) was used for detailed structural comparison with the GlcNAc-PDase model (30).

Both proteins possess an overall β-sandwich-like fold comprised of β-α-β motifs. The GlcNAc-PDase structural region aligns to *Bc*-SMase with a Cα RMSD of ∼4.0 Å indicating global structural homology (Fig. 5B). As EEP superfamily phosphodiesterase enzymes, both GlcNAc-PDase and *Bc*-SMase activity are dependent on divalent metal ion binding. In the *Bc*-SMase crystal structure, two Co^2+^ ions are coordinated by 6 residues in a conserved cleft that has been proposed to directly position the substrate scissile phosphodiester bond for hydrolytic cleavage (30). The *Bc*-SMase metal binding residues are in near-identical positions with homologous residues of GlcNAc-PDase implying the latter also coordinates two metal ions for correct scissile bond orientation (Fig. 5C). Indeed, single site-directed substitutions to alanine for these residues significantly diminish the phosphodiesterase activity of GlcNAc-PDase on 4MU-ý-GlcNAc-6-PC, except for Glu89 (Fig. 5D). Predicted structures of the three tested homologous enzymes selected from the Clostridia group sequence cluster (Table 2) also possess structural conservation for this metal ion binding site (Fig. S14).

### Detecting GlcNAc-6-PC in parasitic nematode glycans

The ability of GlcNAc-PDase to remove PC from GlcNAc-6-PC from intact glycans was investigated. Recent characterization of the glycome of the filarial parasitic nematode *B. malayi* revealed the presence of glycans bearing zwitterionic sugar modifications on both GlcNAc and mannose (Man-PC) (9). In this study, GlcNAc-6-PC was observed in the terminal position of N-glycan outer-arms and in glycosphingolipids (GSLs). Thus, we explored if GlcNAc-PDase could remove PC from GlcNAc-6-PC in intact *B. malayi* glycans. Four fractions containing glycans obtained from *B. malayi* were used as substrates for digestion with IVTT-produced GlcNAc-PDase. Reaction products were analyzed by MALDI-TOF mass spectrometry.

Our *B. malayi* N-glycan substrate has a MALDI ion profile consisting of eight previously characterized N-glycans species (Fig. 6A). Of these, four N-glycans contain PC-modified mannose (structures 1, 4, 6, and 7) and one has a terminal GlcNAc-6-PC (structure 3). Following GlcNAc-PDase digestion, only the ion signature of structure 3 changed (from *m/z* 1398.335 [M-H]^-^ to *m/z* 1233.351 [M-H]^-^) consistent with loss of a single PC (Δ165 Da) from its terminal GlcNAc. No *m/z* changes were observed for any of the Man-PC structures. Thus, GlcNAc-PDase was able to access and remove PC from intact N-glycans and shows the same GlcNAc selectivity observed in our monosaccharide experiments (above).

The first *B. malayi* GSL glycan substrate contains six different structures, five of which harbor GlcNAc-6-PC (Fig. 6B; structures 1, 3, 4, 5 and 6). Upon digestion with GlcNAc-PDase, the m/z values of structures 1 and 5 each shifted by 165 Da due to the loss of PC. Interestingly, the signatures of GlcNAc-6-PC-containing structures 3, 4, and 6 were unaltered. These structures each have PC esterified to a GlcNAc that is internal to the glycan, suggesting that GlcNAc-6-PC must be in a terminal position for PC to be removed by GlcNAc-PDase. It is also noteworthy that only partial digestion of structure 5 was observed, suggesting this glycan fraction comprises a mixture of isomeric structures for which not all GlcNAc-6-PC is in the terminal position.

We further assessed the ability of GlcNAc-PDase to be used in conjunction with an exoglycosidase to verify the presence of internal GlcNAc-6-PC in a GSL glycan. In this experiment, a second GSL glycan substrate having three GlcNAc-6-PC-containing structures was used (Fig. S15A). Treatment with β-*N*-Acetylhexosaminidasef resulted in removal of unsubstituted GalNAc and GlcNAc residues from structures 1 and 3, respectively, to yield structure 4 (*m/z* 830.276 [M-H]^-^) that possesses terminal GlcNAc-6-PC (Fig. S15B.i). Structure 4 became further susceptible to digestion with hexosaminidase only after treatment with GlcNAc-PDase (Fig. S15B.ii). These data again support the idea that GlcNAc-6-PC must be in a terminal position for PC to be removed by GlcNAc-PDase. A similar experiment was conducted with an N-glycan substrate that bears terminal Man-PC (*m/z* 1033.365 [M-H]^-^) (Fig. S16B.i). No digestion of this glycan was observed after sequential treatment with α1-2,3,6 Mannosidase and GlcNAc-PDase (Fig. S16B.ii), confirming that GlcNAc-PDase does not act upon Man-PC, either in a terminal or internal position. Together, these data illustrate the analytical application of GlcNAc-PDase with and without exoglycosidases to verify the presence and location of GlcNAc-6-PC within a glycan.

## DISCUSSION

In the present study, we used functional metagenomic screening to identify active phosphodiesterases produced by human gut microbes that remove zwitterions from carbon-6 of GlcNAc. We describe discovery of a new family of sugar-specific phosphodiesterases having a unique specificity and that functionally define a previously uncharacterized sequence cluster within the large and diverse EEP protein superfamily. We characterized the specificity and biochemical requirements of several members of this protein subfamily and demonstrate an analytical application of this specificity to confirm the presence of GlcNAc-6-PC on glycans isolated from the parasitic nematode *B. malayi*.

Despite the wide distribution of zwitterionic modifications of glycans and glycolipids, few phosphodiesterases that specifically remove these groups from sugars have been described. In yeast and mammalian GPI anchoring, proteins have been implicated in hydrolysis of side-branching PE residues from the GPI glycan. A recombinant version of the mammalian protein PGAP5 can remove PE from the second GPI mannose (Man-2) *in vitro* (21). Yeast proteins with similarity to PGAP5 (Ted1p and Cdc1p), have also been genetically implicated in removal of PE from the innermost two mannoses of the GPI glycan (22, 31). All three proteins share some sequence similarity (22) and each are members of the MMPE1 metallophosphatase family (cd081165). In a second example, a phosphorylcholine esterase (PCE) from *Streptococcus pneumoniae* removes PC from *N*-acetylgalactosamine (GalNAc) in pneumococcal teichoic and lipoteichoic acid (23). PCE is a member of the MBL-fold metallohydrolase (cd07731) family. None of the aforementioned proteins share sequence similarity with GlcNAc-PDase, and none are members of the EEP superfamily.

An interesting biochemical property of GlcNAc-PDase and its close homologs is their high degree of sugar-selectivity. Assays performed using several synthetic zwitterion-modified monosaccharide substrates showed that GlcNAc-PDase has a marked preference for GlcNAc-6-PC or -PE substrates with significantly weaker activity on Glc-6-PC or -PE. This suggests that 2-aminoacetylation of the glucose ring significantly benefits efficient catalysis. Additionally, mannose (a C2 epimer of glucose) and galactose (a C4 epimer of glucose) bearing zwitterions at C6 were not recognized as substrates further suggesting that GlcNAc-PDase likely discriminates its substrate through interaction with hydroxyls at C2 and C4 of GlcNAc. The notion of sugar-selectivity is further supported by the inability of GlcNAc-PDase to hydrolyze the sugarless substrate 4-methylumbelliferyl-PC (4MU-PC) indicating that the presence of a PC phosphodiester linkage alone is not sufficient to permit catalysis. Together, these observations support the conclusion that GlcNAc-PDase and the related proteins in the Clostridia group are sugar-specific phosphodiesterases with remarkable selectivity for GlcNAc-6-PC or -PE.

Recent advances in protein structure prediction allowed for comparative analysis of high-confidence models of GlcNAc-PDase with the *Bc*-SMase-Co^2+^ complex crystal structure (32, 33). GlcNAc-PDase and selected homologs possess structurally conserved EEP metal-binding residues with *Bc*-SMase that likely coordinate divalent metal ions in the active site for hydrolytic activity (30) as supported by mutagenesis. However, determining the structural basis for zwitterion recognition and the sugar-selectivity displayed by this enzyme family will require experimental structure determination in complex with metal ions and substrate molecules.

Analysis of glycans containing zwitterions has historically been challenging. Depending on a variety of technical factors (*e.g*., ionization mode, instrument resolution), mass spectrometry alone can easily miss the presence of a zwitterion on a glycan or fail to discriminate it from other modifications like sulfate (nicely discussed in reference 3). Evidence of PC or PE presence on a glycan can be bolstered by chemical removal of phosphate esters using hydrofluoric acid treatment. However, this can also remove other sugars like fucose and galactofuranose and is not applicable to experiments involving living cells. Thus, sugar-specific phosphodiesterases, used in conjunction with additional exoglycosidases, represent compelling tools for glycan sequencing workflows to confirm the presence of zwitterions. The demonstrated ability of GlcNAc-PDase to hydrolyze PC from GlcNAc-6-PC of intact N-glycans and GSLs suggests its potential for *in situ* analysis of parasite cell surfaces, where GlcNAc is a prevalent modification, and better definition of glycans at the host-pathogen interface.

## EXPERIMENTAL PROCEDURES

*Chemicals, reagents, and enzymes* β-*N*-Acetylhexosaminidasef, RNase inhibitor murine, Q5^®^ Hot Start High-Fidelity 2X Master Mix, endoglycoceramidase I (EGCase I), NEB Express *I^q^* Competent *E. coli, N*-Glycosidase F (PNGase F) (glycerol-free, recombinant), α1-2,3,6 Mannosidase, and GlycoBuffer 1 were from New England Biolabs (Ipswich, MA, USA). 4-Methylumbelliferyl *N*-acetyl-β-D-glucosaminide-6-phosphorylcholine (4MU-ý-GlcNAc-6-PC) and 4-Methylumbelliferyl phosphorylcholine (4MU-PC) were from Biosynth (Staad, Switzerland). 4-Methylumbelliferyl *N*-acetyl-β-D-glucosaminide (4MU-ý-GlcNAc) was from Dextra Laboratories Ltd (Reading, UK). *N*-Acetyl-D-glucosamine-6-phosphorylcholine (GlcNAc-6-PC) and *N*-acetyl-D-glucosamine-6-phosphoethanolamine (GlcNAc-6-PE) were synthesized by Creative Biolabs (Shirley, NY, USA). 6-O-phosphorylcholine-D-glucopyranose (Glc-6-PC), 6-O-phosphorylcholine-D-galactopyranose (Gal-6-PC), 6-O-phosphorylcholine-D-mannopyranose (Man-6-PC), 6-O-aminoethylphosphonato-D-glucopyranose (Glc-6-PE), 6-O-aminoethylphosphonato-D-mannopyranose (Man-6-PE), and 2-O-aminoethylphosphonato-D-mannopyranose (Man-2-PE) were synthesized by GlycoUniverse (Potsdam, Germany). Y-PER™ Yeast Protein Extraction Reagent was from Thermo Fisher Scientific Inc. (Waltham, MA, USA). The CopyControl™ Fosmid autoinduction solution was from Lucigen Corporation (Middleton, WI, USA). The Bruker^®^ Peptide Calibration Standard and 2,5-dihydroxybenzoic acid (DHB) was from Bruker Daltonics (Billerica, MA, USA).

### Construction of a human gut microbiome metagenomic library

A fosmid metagenomic library was constructed using DNA extracted from a human gut microbiome fecal sample as previously described (24). Briefly, DNA was isolated from 100 mg of fecal sample. Isolated DNA was used to construct a fosmid library using the CopyControl™ Fosmid Library Production kit (Lucigen Corporation) following the manufacturer’s instructions.

### Phosphodiesterase screening from a human gut microbiome metagenomic library

A total of 6,144 fosmid clones from a human gut microbiome library (24) were screened for GlcNAc-6-PC phosphodiesterase activity in a 384 well plate format. Library clones were cultured overnight at 37 °C in 50 µL LB containing 12.5 µg/mL chloramphenicol and 1X CopyControl™ Fosmid autoinduction solution. Cultures were lysed by addition of 50 µL of Y-PER supplemented with 40 µg/mL of 4MU-β-GlcNAc-6-PC and 1 U/mL of β-*N-* Acetylhexosaminidasef per well. Reactions were incubated at 37 °C for 48 h. Fluorescence was read at λex = 365 nm and λem = 445 nm using a SpectraMax Plus 384 Microplate reader (Molecular Devices, San Rose, CA, USA) at different timepoints (1, 5, 24, 48 h). Hits were defined as RFU values above 3 standard deviations from the mean. Hits from the primary screen were validated by repeating the screening protocol. The hit definition was then refined to wells with RFU values that were 10 standard deviations above the mean in at least two timepoints.

### Fosmid sequencing

Fosmids from each hit were isolated by BioS&T (Quebec, Canada) or using the FosmidMAX™ DNA Purification Kit (Lucigen Corporation). Isolated fosmids were prepared for Illumina sequencing using the plexWell™ 96 kit (seqWell, Danvers, MA, USA) following the manufacturer’s instructions. Sequence assembly was performed by seqWell. Open reading frames were predicted using MetaGeneMark (34).

### *In vitro* expression of enzyme candidates and mutants

Interesting ORF candidates were selected for *in vitro* expression based on their predicted annotation from a BLASTP search. Selected candidates were expressed *in vitro* using the PURExpress^®^ *In Vitro* Protein Synthesis Kit (New England Biolabs) following the manufacturer’s instructions. Briefly, DNA templates were prepared by PCR using Q5^®^ Hot Start High-Fidelity 2X Master Mix in a 60 µL total reaction volume. Primers used are in Table S1. PCR products were purified using the Monarch^®^ PCR & DNA clean-up kit (New England Biolabs). *In vitro* expression was performed for 2 h at 37 °C. A positive control PURExpress^®^ reaction was performed in 25 µL total reaction volume using 2 µL of 125 ng/µL dihydrofolate receptor (DHFR)-expressed plasmid supplied with the kit. Expressed proteins were separated on a Novex 10-20% Tris-glycine gel (Thermo Fisher Scientific Inc.) using 2.5 µL of the reaction. PURExpress^®^ reactions were then assayed for phosphodiesterase activity by mixing 7 µL of PURExpress^®^ mixtures with 2 µL of 100 µg/mL 4MU-Β-GlcNAc-6-PC, and 5 units of β-*N*-Acetylhexosaminidasef. To assess hexosaminidase activity expressed from fosmid clones, reactions were assayed as indicated above, but in the absence of exogenous β-*N*-Acetylhexosaminidasef. Reactions were incubated for 1 h at 37 °C and fluorescence was read at λex = 365 nm and λem = 445 nm using a SpectraMax Plus 384 microplate reader.

Six GlcNAc-PDase mutants (N60A, E89A, D204A, N206A, D266A, and H267A) were synthesized with *E. coli* codon optimization, and cloned into pUC57 (GenScript, Piscataway, NJ).

Mutants were expressed *in vitro* as described above. Activity of each mutant was assayed on 4MU-ý-GlcNAc-6-PC in triplicate as described above for wild-type GlcNAc-PDase.

### *In vivo* expression and purification of recombinant GlcNAc-PDase

DNA encoding GlcNAc-PDase with a C-terminal 8-His tag was codon optimized for *E. coli* and synthesized (Genscript). GlcNAc-PDase-8His was subcloned into expression vector pJS119k (35). Luria–Bertani (LB) medium supplemented with 25 μg/mL kanamycin was inoculated with NEB Express *I^q^* Competent *E. coli* cells carrying the pJS119k-GlcNAc-PDase-8His plasmid and grown at 37 °C until the OD600 reached 0.4. Isopropyl β-D-thiogalactoside (IPTG) was then added to induce expression and the culture was incubated at 16 °C overnight with shaking. Cells were harvested by centrifugation, re-suspended in column buffer A (50 mM Tris-HCl, pH 7.5, 300 mM NaCl and 50 mM imidazole), and lysed by passing twice through a Shear Jet HL60 homogenizer (Dihydromatics, Maynard, MA) at 22 KPsi. The lysate was loaded onto a HisTrap FF column (Cytiva Life Sciences, Uppsala, Sweden) that had been pre-equilibrated with column buffer A. The column was washed with column buffer A and bound protein was eluted with column buffer B (50 mM Tris-HCl, pH 7.5, 300 mM NaCl and 1M imidazole). Fractions containing purified recombinant F4-GlcNAc-PDase-8His were pooled and diluted 10-fold with column buffer A and re-loaded onto a HisTrap FF column that had been pre-equilibrated with column buffer A. Bound protein was again eluted with column buffer B.

### GlcNAc-PDase biochemical characterization

To assess the metal ion requirement of the enzyme, 0.2 U of GlcNAc-PDase was incubated with 200 ng of 4MU-ý-GlcNAc-6-PC, 5 U of β-*N*-Acetylhexosaminidasef in 20 mM MES, pH 6.5, with 5 mM of MgCl2, CaCl2, CuSO4, NiCl2, MnSO4, FeSO4, CoCl2, or ZnSO4 for 1 h and 19 h at 37 °C. Reactions were performed in triplicate. Fluorescence was read at λex = 365 nm and λem = 445 nm using a SpectraMax Plus 384 microplate reader.

To determine the optimal pH, 0.3 U of GlcNAc-PDase was incubated for 1 h at 37 °C with 1 mM GlcNAc-6-PC, 5 mM MgCl2, and buffers ranging from pH 4.5 to 10.8 (from pH 4.5 to 5.5, 50 mM sodium acetate; from pH 6.0 to 7.0, 20 mM sodium phosphate; from pH 7.5 to 9.0, 50 mM Tris-HCl; pH 9.9, 20 mM CAPS; and pH 10.8, 50 mM carbonate buffer) in a 10 µL final reaction volume. Reactions were performed in triplicate. After 1 h incubation at 37 °C, 5 µL of each reaction was dried using a centrifuge concentrator (Vacufuge plus, Eppendorf, Hamburg, Germany) for 30 min. Each dried reaction was procainamide labeled. Procainamide solution was prepared by first dissolving 12 mg of procainamide in 110 µL of 70% dimethyl sulfoxide (DMSO)/ 30% acetic acid solution. This mixture and 25 µL of water were then transferred to a separate tube containing 6 mg of sodium cyanoborohydride. Dried GlcNAc-PDase reactions were incubated for 1 h at 65 °C with 20 µL freshly made procainamide solution. A 1 µL aliquot was diluted in 9 µL of 100% ACN and separated by UPLC (see below).

The optimal temperature of GlcNAc-PDase catalysis was determined by incubating 0.3 U GlcNAc-PDase with 1 mM GlcNAc-6-PC, 5 mM MgCl2, and 50 mM Tris-HCl pH 8.0 for 1 h at 8 temperatures ranging from 15 to 70 °C. Reactions were performed in triplicate, dried, and procainamide-labeled as previously described. A 1 µL aliquot was diluted in 9 µL of 100% acetonitrile and separated by UPLC (see below).

### Ultra-Performance Liquid Chromatography and Mass Spectrometry

Procainamide-labeled samples were separated by ultra-performance liquid chromatography (UPLC) using a ACQUITY UPLC glycan BEH amide column 130Å (2.1 × 150 mm, 1.7 μm) from Waters on a H-Class ACQUITY instrument (Waters, Milford, MA). Solvent A was 50 mM ammonium formate buffer, pH 4.4 and solvent B was acetonitrile (ACN). The gradient used with 130Å (2.1 × 150 mm, 1.7 μm) column was 0–35 min, 12–47% solvent A; 35-35.5 min, 47-70% solvent A; 35.5-36.0 min, 70% solvent A; 36-36.5 min, 70-12% solvent A; 36.5-40 min, 12% solvent A. The flow rate was 0.4 mL/min. The injection volume was 3 μL and the sample was prepared in 100% (v/v) ACN. Samples were kept at 5 °C prior to injection and the separation temperature was 40 °C. The fluorescence detection wavelengths for procainamide were λex= 308 nm and λem= 359 nm, with a data collection rate of 20 Hz.

Conditions for inline mass detection using the ACQUITY quadrupole QDa (Waters) were as follows: Electrospray ionization (ESI) in positive mode; capillary voltage, 1.5 kV; cone voltage, 15 V; sampling frequency, 5 Hz; probe temperature 400 °C. The QDa analysis was performed using full scan mode, and the mass range was set at m/z 50–800. Single ion recording (SIR) mode was used as well to monitor individual glycans. Waters Empower 3 chromatography workstation software was used for data processing including traditional integration algorithm, no smoothing of the spectra and manual peak picking.

### Family analysis of GlcNAc-PDase

Protein sequences of EEP superfamily members were obtained from the following sources: [1] UniProt entries for mouse PGAP2IP (Q91YL7), *S. cerevisiae* Cwh43p (P25618), and *S. pombe* Cwh43p (Q9HDZ2); [2] BLASTP hits from bacteria, archaea, and eukarya using GlcNAc-PDase and Q91YL7 as queries to the NCBI nr database were separately collected until the sets of hits to the two queries overlapped, after which duplicates were removed; [3] BLASTP hits to WP_159773962 (jacalin-related lectin EEP domain) from bacteria, archaea, and eukarya (36); [4] representative proteins from each domain in NCBI’s Conserved Domain Database (CDD) that is a subset of cd08372 (EEP domain).

Protein sequences from the sources above were combined, and redundancy was reduced using CD-Hit (37) to replace sets of sequences sharing >70% identity with a single representative. Sequences that appeared to be incomplete, either based on annotation or on multiple sequence alignment, were removed. The final set comprised 900 sequences.

Sequences were clustered in two dimensions using CLANS (38) run under the MPI Bioinformatics Toolkit (39). BLAST connections with P-values < 1e-04 are shown as edges, and the clustering was run to convergence (typically 10,000-30,000 iterations). Sequences from the GlcNAc-PDase cluster were aligned with MUSCLE v. 5.1 (40) (Fig. S4) within the Geneious software package, regions outside the EEP domain (which in the case of GlcNAc-PDase comprised N-terminal residues 1-53) were removed, and the 186 GlcNAc-PDase EEP domain sequences were re-clustered using the same procedure.

### Specificity of GlcNAc-PDase and related proteins

Three closely related gene sequences to GlcNAc-PDase (NLD61619, MBE6669927, and HCG67795) were identified from the Clostridia group of the GlcNAc-PDase family analysis (see above). Signal peptide sequences predicted using SignalP 5.0 (41) were removed from each gene sequence. Gene constructs were synthesized, *E. coli* optimized, and cloned into pUC57 (GenScript). Each sequence was expressed *in vitro* using the PURExpress^®^ *In Vitro* Protein Synthesis Kit as described above (Fig. S5A). Activity of each protein was assayed on 4MU-ý-GlcNAc-6-PC as described for GlcNAc-PDase (Fig. S5B).

The specificity of GlcNAc-PDase and related proteins was assessed by incubating purified or *in vitro* expressed material with the following custom synthesized monosaccharides: GlcNAc-6-PC, Glc-6-PC, Gal-6-PC, Man-6-PC, GlcNAc-6-PE, Glc-6-PE, Man-6-PE, and Man-2-PE. Reactions were performed in triplicate using 1 U of purified GlcNAc-PDase or 1 µL of *in vitro* expressed material, 10 nmol of each monosaccharide, 50 mM Tris-HCl pH 8.0, and 5 mM MgCl2 in a final volume of 10 µL. After 1 h incubation at 37 °C, 5 µL of each reaction was dried using a Vacufuge plus for 30 min. Dried reactions were procainamide labeled and resolved on the UPLC as described above.

### Structural analysis of GlcNAc-PDase and related proteins

Structural predictions of GlcNAc-PDase and the three homologs described above were performed using the Colabfold platform (33) with AlphaFold2 (32) and substituted homology detection and sequence alignment pairing with MMSeq2 (42) and HHsearch (43). Multiple sequence alignment (MSA) coverage and pLDDT plots are provided in Supporting Information Fig. S17. PyMol was used for visualizations and interpretation (44). The termini, that include the secretion signal sequence, were predicted with low confidence. Superposition of the GlcNAc-PDase structured region (residues 50-262) with the *Bc-*SMase-Co^2+^ complex (PDB ID 2DDS) was performed for Cα atoms using the *align* command with default parameters for an RMSD of 3.9 Å (157 atoms).

### Extraction and release of *B. malayi* N-linked and GSL glycans

Purified glycans from the parasitic filarial nematode *B. malayi* were generated in a previous study (9). In summary, parasite glycoproteins and glycolipids from approximately 600 adult female worms were isolated by tissue homogenization in MQ (MilliQ water), methanol (MeOH) and chloroform in a final 4:7:13 ratio of MQ:MeOH:chloroform. Samples were sonicated and centrifuged at 4000 rpm for 5 minutes and the upper phase was removed and replaced by 50% MeOH. This step was repeated twice and (glyco)lipids from the upper phase of the extraction were subsequently purified using octadecylsilane (C18) cartridges (BAKERBOND^®^ spe™, JT Baker^®^, Avantor, Radnor Township, PA, USA) as described elsewhere (45). Glycolipids eluted from cartridges were dried under nitrogen (N2) flow and reconstituted in 500 µl 50 mM sodium acetate (NaAc) pH 5.0 containing 0.1% natrium taurodeoxycholate. To release glycosphingolipid (GSL) derived glycans, lipid extracts were treated with a total of 32 mU of recombinant EGCase I over 48 h at 37 °C. An aliquot (16 mU) of the enzyme was first added while another 16 mU of EGCase I was added after 24h of incubation. Simultaneously, (glyco-)proteins in the lower phase from the protein/lipid separation procedure described above were pelleted using excess volumes of MeOH. Protein pellets were dried under N2 flow and then homogenized in PBS with 1.3% SDS and 0.1% β-mercaptoethanol. Denaturation was performed for 10 min at 95 °C, samples were cooled to room temperature and Nonidet P-40 (1.3% final concentration) was added to each sample. N-linked glycans were released using 3500 units of PNGase F and incubation at 37 °C for 24 h.

Following their enzymatic release, N-linked and GSL glycans were cleaned-up sequentially on C18 and carbon cartridges (Supelclean™ ENVI-Carb SPE, MilliporeSigma, Burlington, MA, USA) according to the protocol previously described (45). Eluted glycans were dried using a Vacufuge plus. The glycans were labelled with 2-aminobenzoic acid (2-AA) by reductive amination with sodium cyanoborohydride as described (46). Labelling was performed for 2h at 65 °C. To remove labelling reagent excess, ACN was added to a final concentration of 75% and the sample loaded onto Bio-Gel P10 Gel resin (Bio-Rad, Hercules, CA, USA) previously conditioned with 80% ACN. Glycans were eluted with MQ and dried using a Vacufuge plus.

### Glycan purification using UPLC fractionation

Released and AA-labeled glycans were next purified using a bidimensional UPLC protocol as part of the glycan microarray construction procedure detailed previously (9). Briefly, glycans were separated using the Dionex UltiMate 3000 system (Thermo Fisher Scientific Inc.), first by normal phase UPLC using hydrophilic interaction chromatography, and then by reverse-phase UPLC on a C18 columns. Fractions were collected, dried down in a Vacufuge plus and re-dissolved in 50 µL of MQ. Glycan content was analyzed by MALDI-TOF-MS as described below. *B. malayi* glycans have been structurally characterized using a combination of orthogonal approaches including glycan sequencing procedures and MS/MS that have been extensively described previously (9). We selected four fractions shown previously to contain zwitterionic glycans representative of the different contexts of PC-substitution identified in *B. malayi* glycans (9).

### Digestion of zwitterionic *B. malayi* glycans with the GlcNAc-PDase

The four fractions containing PC-substituted glycans were subjected to digestion with GlcNAc-PDase alone or in combination with another exoglycosidase, either β-*N*-Acetylhexosaminidasef or the α1-2,3,6 Mannosidase. Digestions were performed by mixing 1-2 µL (out of 50 µL of the glycan fractions) with 1 µL of GlycoBuffer 1 (5 mM CaCl2, 50 mM sodium acetate, pH 5.5 at 25 °C) and with 0.5 µL of the PURExpress^®^ reaction mixture containing the *in vitro* expressed GlcNAc-PDase. For selected fractions, 2 µL of either β-*N*-Acetylhexosaminidasef or α1-2,3,6 Mannosidase were additionally used. Reaction volumes were then adjusted to 10 µL total with water and digestions were performed overnight at 37 °C. Negative controls including all the listed reagents minus the glycosidase(s) were run.

Enzyme removal was subsequently performed for all digestion reactions and controls using C18 Millipore^®^ Zip-Tips (MilliporeSigma) as detailed previously (47). As described, glycans were directly eluted onto the MALDI plate in 50% ACN, 0.1% TFA mixed with 10 mg/ml DHB at the end of the clean-up procedure, and were subsequently analyzed using MALDI-TOF-MS.

### MALDI-TOF-MS analysis

MALDI-TOF-MS analysis was performed using an UltrafleXtreme^®^ mass spectrometer (Bruker Daltonics) equipped with 1 kHz Smartbeam II laser technology and controlled by the software FlexControl 3.4 Build 119 (Bruker Daltonics). Samples eluted from Zip-Tips were spotted on a 384 well steel polished target plate in DHB matrix as mentioned above. All spectra were obtained in the negative-ion reflectron mode using Bruker^®^ peptide calibration standard mix for external calibration. Spectra were obtained over a mass window of m/z 700 – 3500 with ion suppression below m/z 700 for a minimum of 20,000 shots (2000 Hz) obtained by manual selection of “sweet spots”. The software FlexAnalysis (Version 3.4, Build 50, Bruker Daltonics) was used for data processing including smoothing of the spectra (Savitzky Golay algorithm, peak width: m/z 0.06, 1 cycle), baseline subtraction (Tophat algorithm) and manual peak picking. Peaks with a signal-to-noise ratio below 5 were excluded as well as known non-glycan peaks such as glucose polymers and contaminants introduced by enzyme treatment. Deprotonated masses of the selected peaks were matched to corresponding glycan structures that were previously elucidated (9).

## Supporting information

This article contains supporting information.

## Data availability

The sequencing data supporting the conclusions of this article are available in the GenBank^™^/EBI Data Bank repository, (OQ439824-OQ439830 in https://www.ncbi.nlm.nih.gov/genbank/). Annotated MALDI-TOF-MS spectra supporting our findings have been made available in the manuscript (main text and supplementary data). In addition, all raw MALDI-TOF-MS data have been deposited in the public repository Glycopost (https://glycopost.glycosmos.org/, project ID: GPST000347).

## Acknowledgments

The authors thank New England Biolabs for basic research funding. We thank Léa Chuzel and Andrew Gardner for helpful comments, advice, and critical review of the manuscript prior to its submission.

## Funding and additional information

This work was supported by New England Biolabs.

