## Supporting Information for "A novel family of sugar-specific phosphodiesterases that remove zwitterionic modifications of *N*-acetylglucosamine"

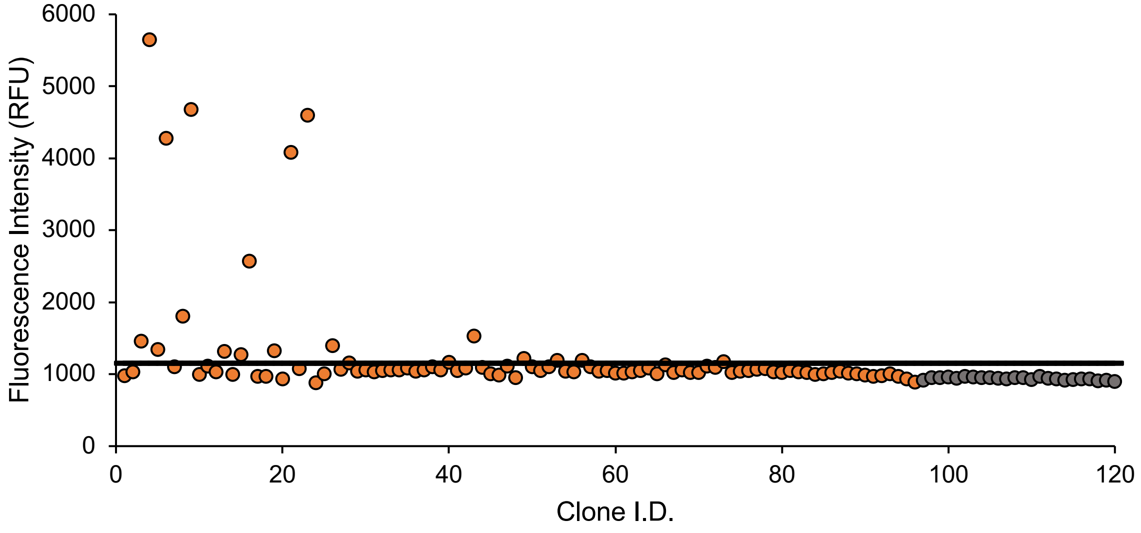


**Figure S1. Functional metagenomic screening.**

A human gut microbiome metagenomic fosmid library was screened with the substrate 4MU-β-GlcNAc-6-PC plus an exogenous hexosaminidase (β-*N*-Acetylhexosaminidase$_f$). Hits from the primary screen were re-screened to determine if the observed activity was reproducible. Shown are fluorescence values at the 24 h timepoint. Metagenomic clones (orange circles) were screened for activity. Replicates of the pCC1 empty fosmid vector (grey circles) were screened as a negative control for background fluorescence. Hits from the re-screen were defined as clones yielding an assay signal at least 10 standard deviations (black line) above the mean background fluorescence.

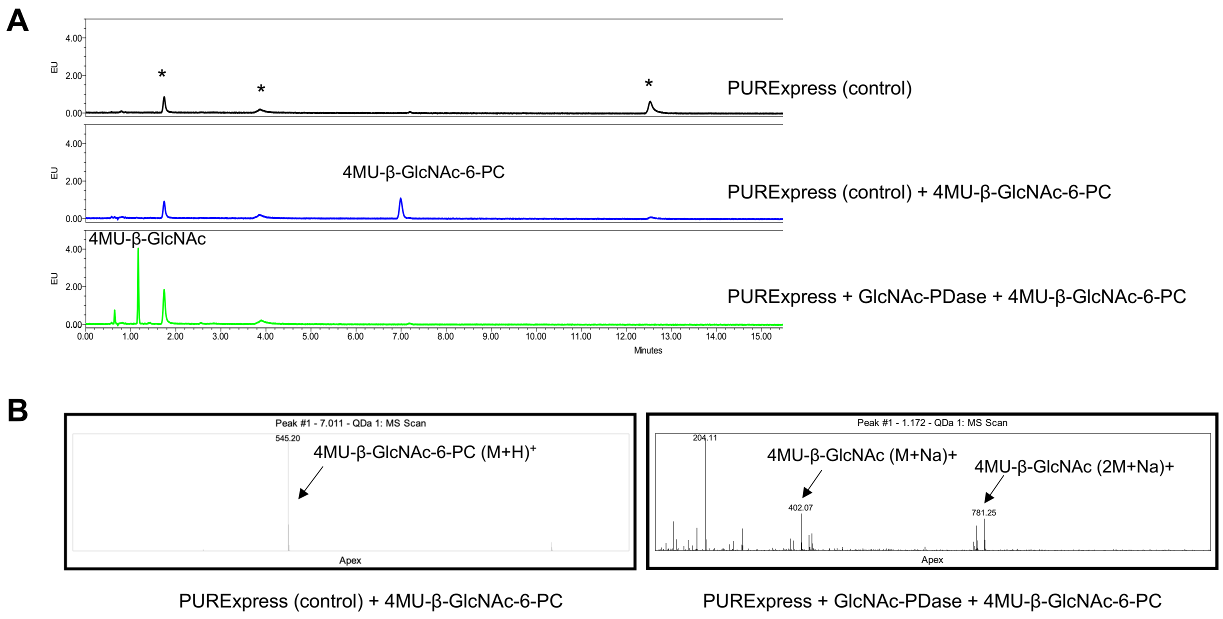


**Figure S2. Activity of GlcNAc-PDase on 4MU-labeled GlcNAc-6-PC substrate analyzed by UPLC-FLR-MS.**

A) UPLC-FLR chromatograms showing separation of the 4MU-β-GlcNAc-6-PC and 4MU-β-GlcNAc before and after incubation with GlcNAc-PDase. Peaks detected due to the intrinsic fluorescence of the PURExpress® components are denoted with an asterisk (*)
B) Analysis of the reaction products from panel A (middle and bottom chromatograms) by UPLC with inline MS.

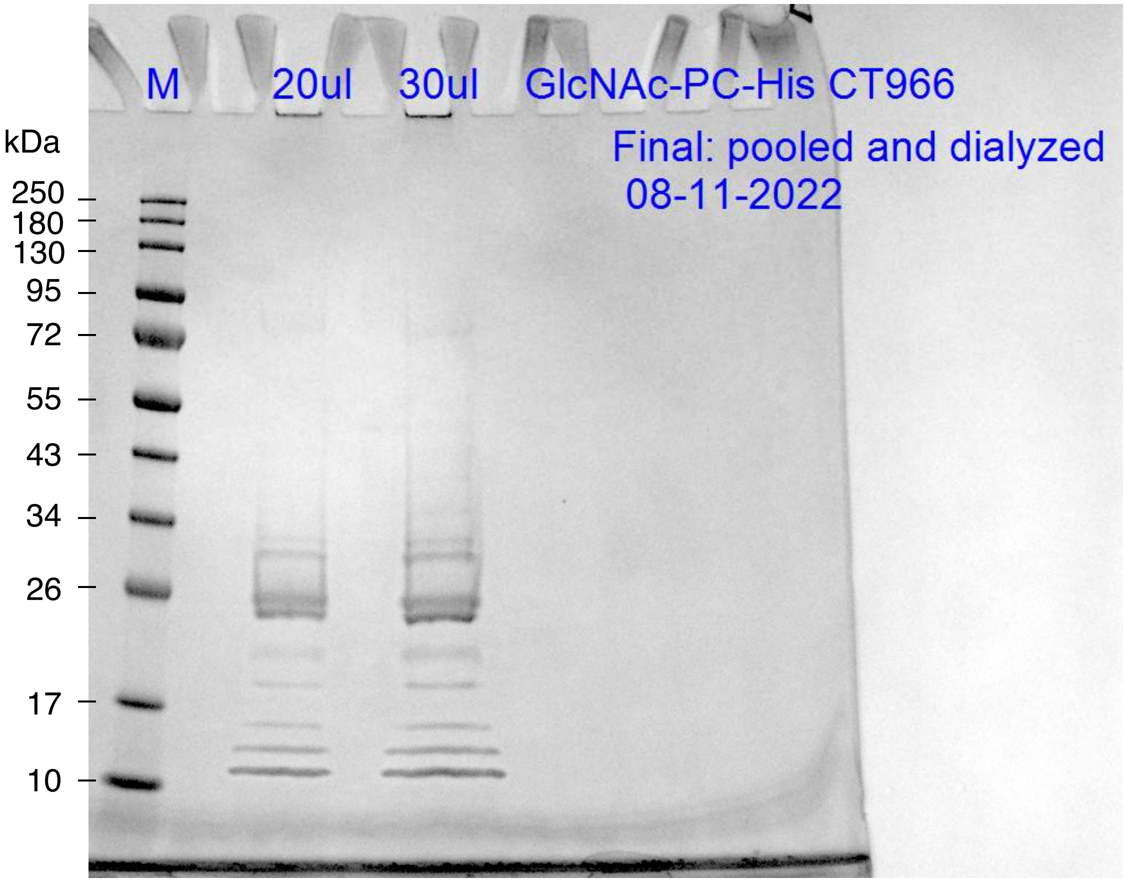


**Figure S3. GlcNAc-PDase-8HIS purification.**

GlcNAc-PDase-8HIS was expressed from the T7 promoter in *E. coli* and purified using Ni-NTA affinity chromatography. The theoretical molecular weight of GlcNAc-PDase is 32 kDa. Purified protein (20μL and 30μL) was loaded on the SDS-PAGE and stained using SimplyBlue™ SafeStain.

```
              1            10        20        30        40        50        60
              |            |         |         |         |         |         |
GlcNAc-PDase  MDMLQ-QIGSALMAGCL-----IVL-QFI-TGMLGFLDGVP---------GSGSKPVAPP
    HCG67795  MKKEYKKLISLILAVAAAGTASALLSACSGSNANTPDISTNSGTVPEEKTVTSPEPETTP
  MBE6669927  MKKLT-ALLTLVLILSL-----TLI-SCI-DEGNISDDQTDGTTLADASTPPAADTTTAY
    NLD61619  MKKISSAVATIAALLAL----------AM---------------------PANAKDETKS

GlcNAc-PDase  ENITPAQ--SVTLRVATYNVKNCVNGT-AVEEIAQDINANDLDIVCLQELDRGGTRAGGK
    HCG67795  EPETTAEPPKLTLRIGSYNIKHGADASGELSLIGGVIKNAGLDIVGVQEVDYKTERVSGV
  MBE6669927  DTTTESQEGLLSLNIGSYNIANGKLVSHDIKVIANDILSKDLDIVGLQEVDKFAKRSKYI
    NLD61619  EMTTASQ--PINIKIASYNILHGAKVNCDYAVLAKDIAEVDPDIIGLQEVDMKTTRIGGV

GlcNAc-PDase  NLLRMLSQKT-LPYYRFFPAIGFPG--------NEYGIGILSRYPFEEVELHRLETGIEE
    HCG67795  DQPAALAAAAGMEYYKFARAIDYRG--------GEYGTLILSRYPIEEFTVTALESGSRE
  MBE6669927  DTMALLSEYTGYKYYHYTVAINIAGNEAVYGQKGEYGTGILSKYPILETKSIKLESGGNE
    NLD61619  DAVEIMAKKAGYKYYRFSKSINLKG--------GGYGTAILSKFPIEEYQTVALESGKHE

GlcNAc-PDase  GRVLGGVTIRADGIPVRIYNTHLSFESMELRTNQITSISETLRGKAPCLLMGDFNLLDFS
    HCG67795  GRSIGHAVINVDGVKVDFFNTHLSYEEKAHRTAQFVKIRELLSDCTTYVLTGDFNTQDYS
  MBE6669927  QRMLGYAKIDVNGQIINFYNTHLSYEDFSIRTGQFETIASLLKDKEYCILTGDFNIAGFG
    NLD61619  KRSVGHAVLRVRGKRLDFFNTHLSYESTKVRAGQFAAIADMTAKCERYIVTGDFNTSDFA

GlcNAc-PDase  ELEGFQGMTPVNTARNPIVTFPCEDGSEYPYLDNILFSAGVECRWVKPYVQTVSDHIMLM
    HCG67795  EFTVLGYGGMLNNAQRHYVTFPGNKSS----IDNIVFSGNFKISKSGTVAESYSDHRMLW
  MBE6669927  EFKSIPFLNTTCNANNWLTTFPSNSSS----IDNILFSNEFALAQSKVLAQGHSDHNMLY
    NLD61619  EFAVFKGATLANNAQHSLPTFPGSSMP----IDNIVLSKGFELISTDIRKNDHSDHYLFY

GlcNAc-PDase  AEVTVANPEVQRDEA
    HCG67795  AELALSI------GD
  MBE6669927  ATLKYR-------SK
    NLD61619  AQTKL--------SE
```

**Figure S4. GlcNAc-PDase and related "Clostridia group" proteins alignment.**

The deduced peptide sequence of GlcNAc-PDase was aligned with three proteins (GenBank numbers: HCG67795, MBE6669927, and NLD61619) from the Clostridia group using the program MUSCLE (https://www.ebi.ac.uk/Tools/msa/muscle/). Identical residues are shaded orange, and gaps introduced into the alignment by the algorithm are denoted with a hyphen.

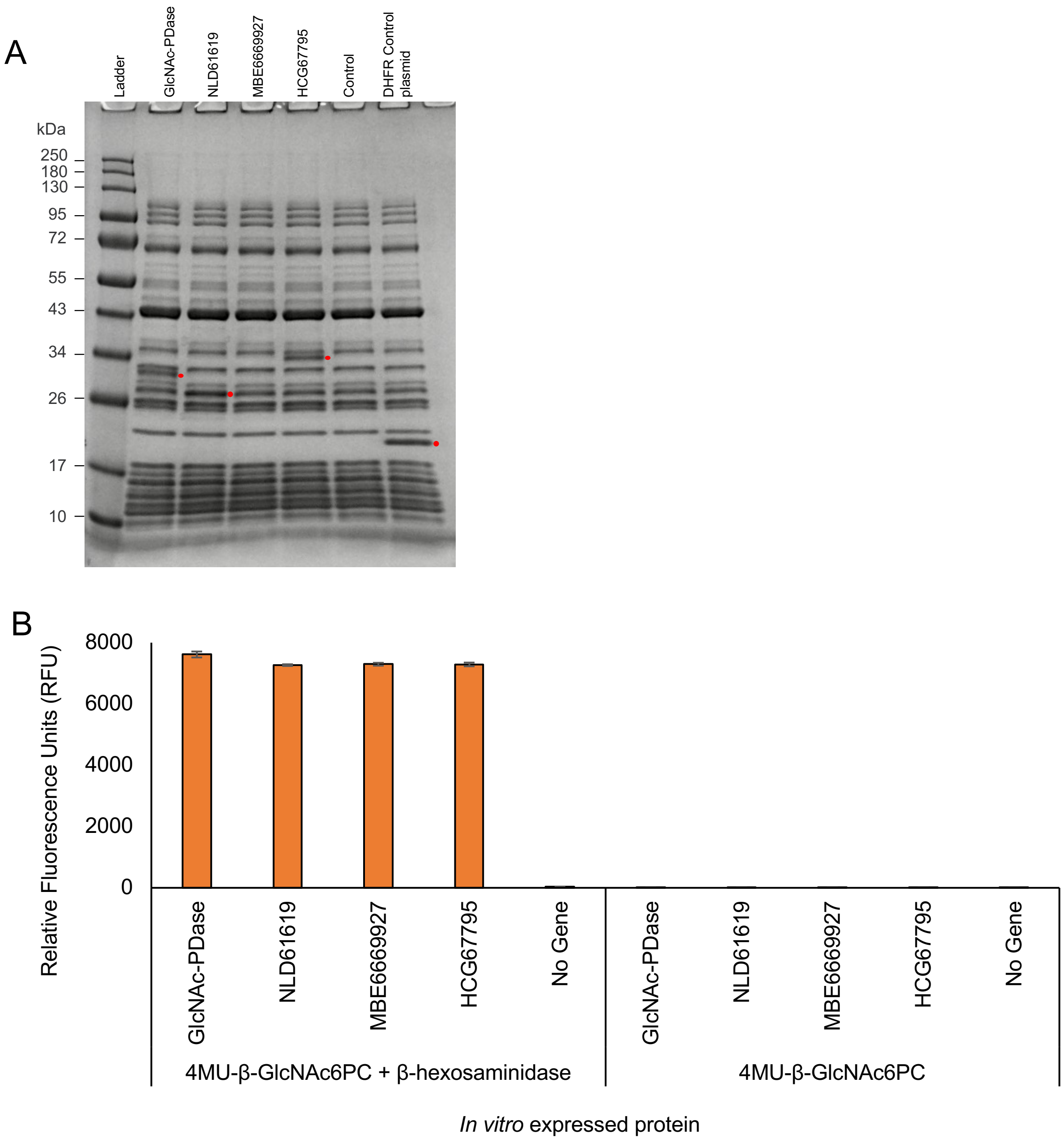


**Figure S5. Expression and activity of "Clostridia group" related proteins.**

A) Three related proteins from the Clostridia group (NLD61619, MBE6669927, and HCG67795) were expressed by *in vitro* transcription and translation (IVTT) using the PURExpress® system (red circles) and compared to GlcNAc-PDase. An empty vector (pCC1) negative control and a PURExpress® DHFR (dihydrofolate reductase) positive control plasmid were also processed.

B) IVTT-produced material was assayed for fluorescence activity on the screening substrate 4MU-β-GlcNAc-6-PC with or without supplemented β-hexosaminidase.

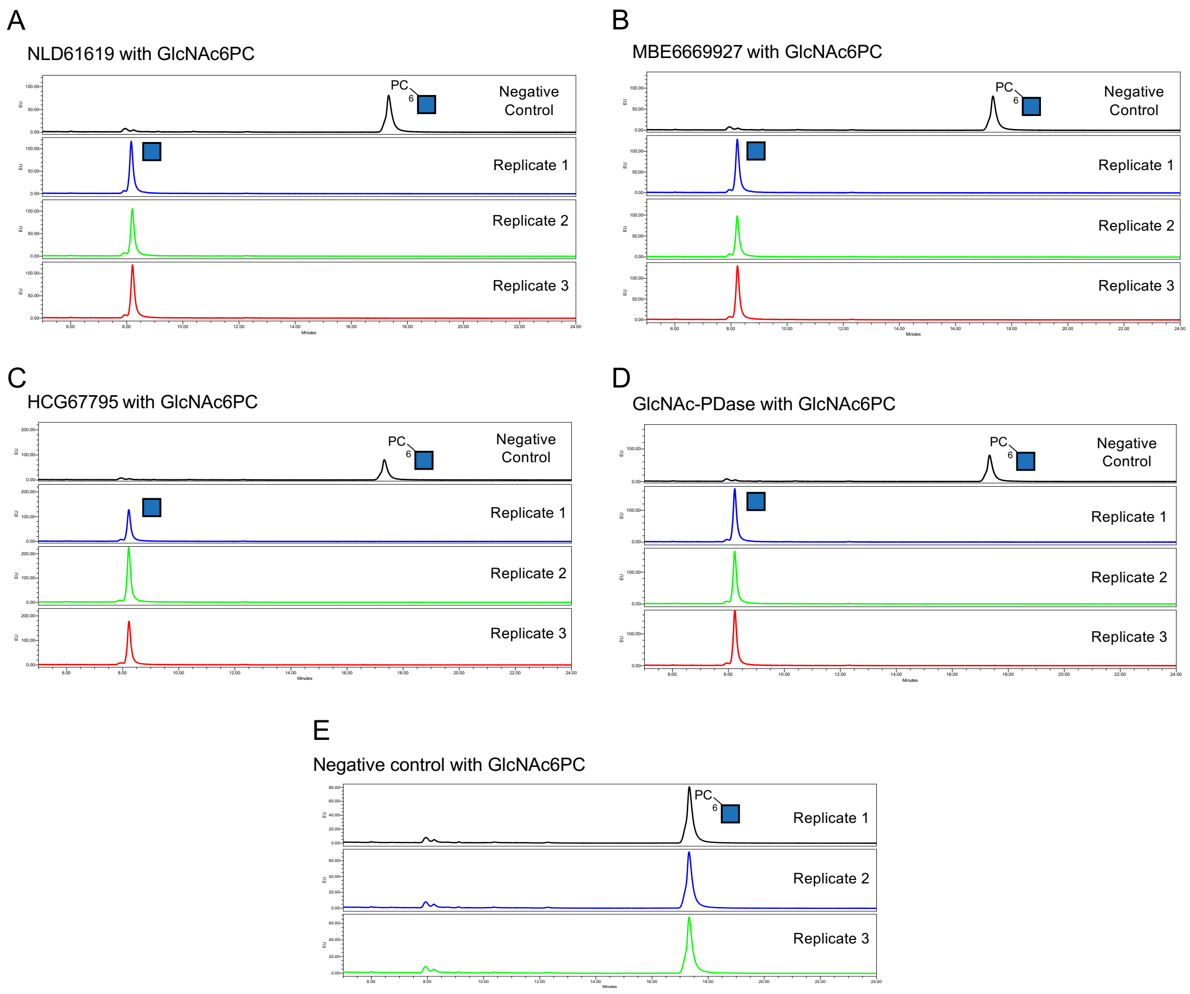


**Figure S6. GlcNAc-PDase and "Clostridia group" protein activity on GlcNAc-6-PC with UPLC-FLR analysis.**

IVTT-produced (A) NLD61619 (related protein 1), (B) MBE6669927 (related protein 2), (C) HCG67795 (related protein 3), (D) GlcNAc-PDase and (E) a negative control were incubated with the monosaccharide *N*-acetyl-D-glucosamine-6-phosphorylcholine (GlcNAc-6-PC) in triplicate. Reactions were procainamide labeled and separated using UPLC-FLR.

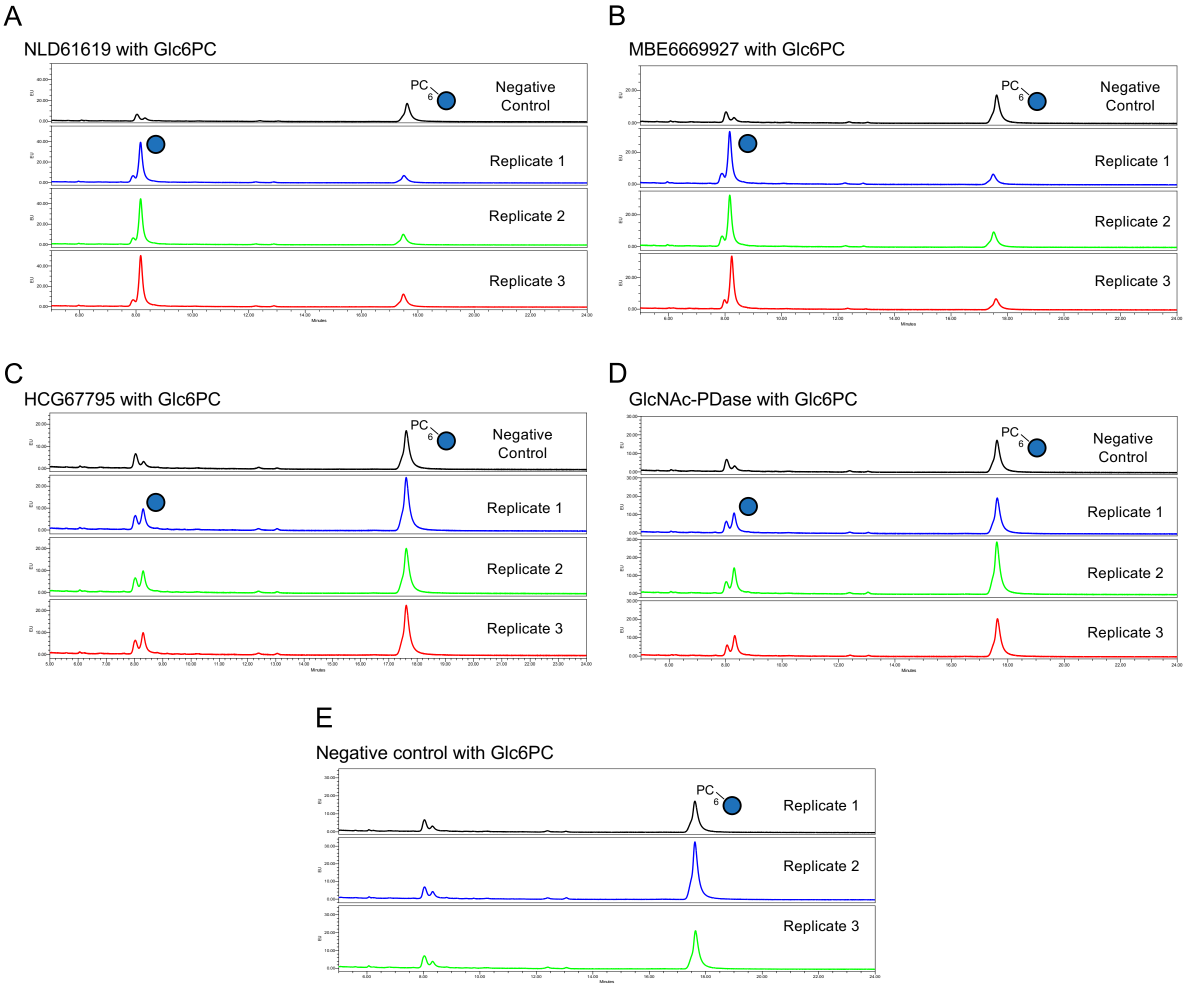


**Figure S7. GlcNAc-PDase and "Clostridia group" protein activity on Glc6PC with UPLC-FLR analysis.**

IVTT-produced (A) NLD61619 (related protein 1), (B) MBE6669927 (related protein 2), (C) HCG67795 (related protein 3), (D) GlcNAc-PDase and (E) a negative control were incubated with the monosaccharide 6-O-phosphorylcholine-D-glucopyranose (Glc-6-PC) in triplicate. Reactions were procainamide labeled and separated using UPLC-FLR.

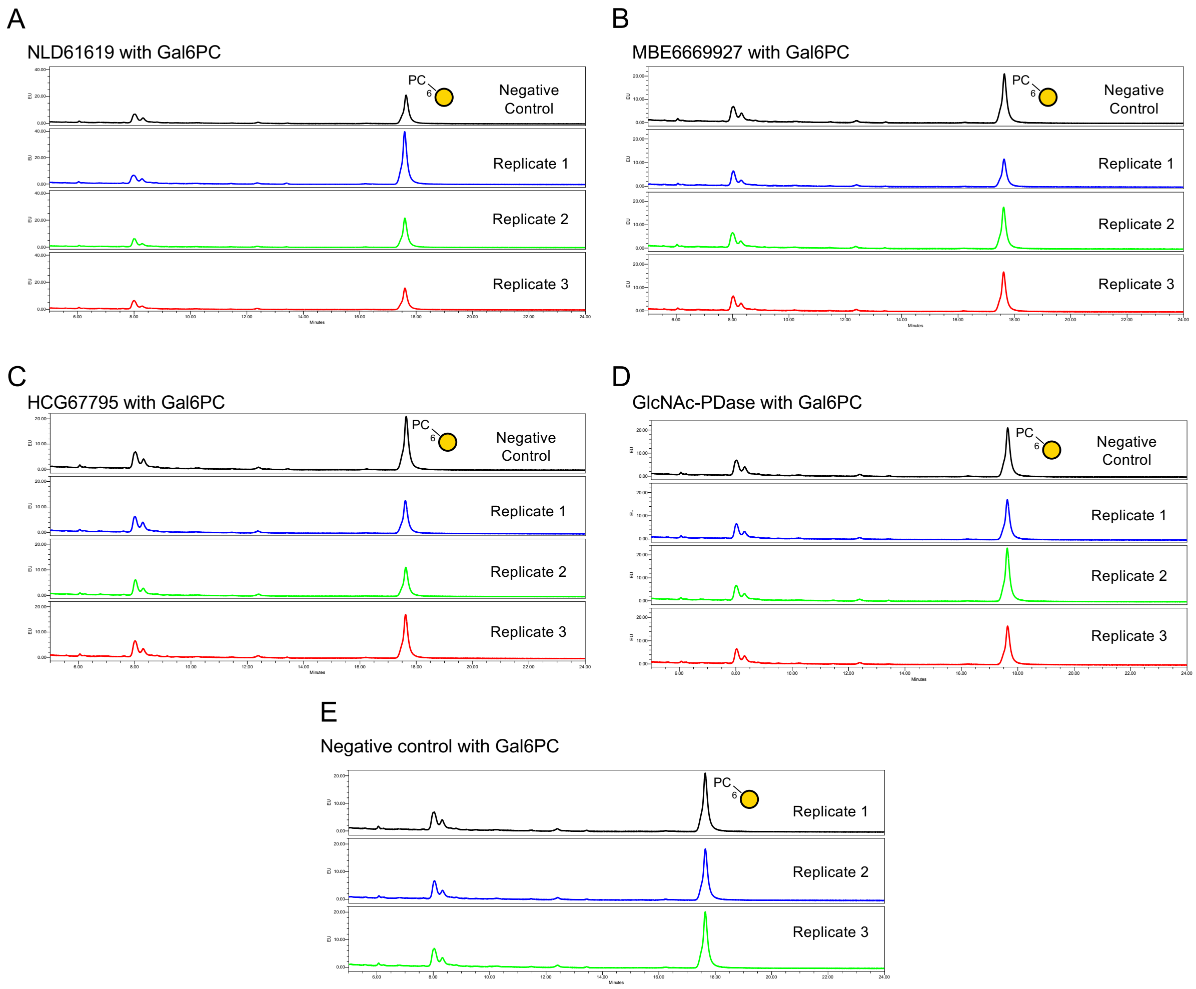


**Figure S8. GlcNAc-PDase and "Clostridia group" protein activity on Gal6PC with UPLC-FLR analysis.**

IVTT-produced (A) NLD61619 (related protein 1), (B) MBE6669927 (related protein 2), (C) HCG67795 (related protein 3), (D) GlcNAc-PDase and (E) a negative control were incubated with the monosaccharide 6-O-phosphorylcholine-D-galactopyranose (Gal-6-PC) in triplicate. Reactions were procainamide labeled and separated using UPLC-FLR.

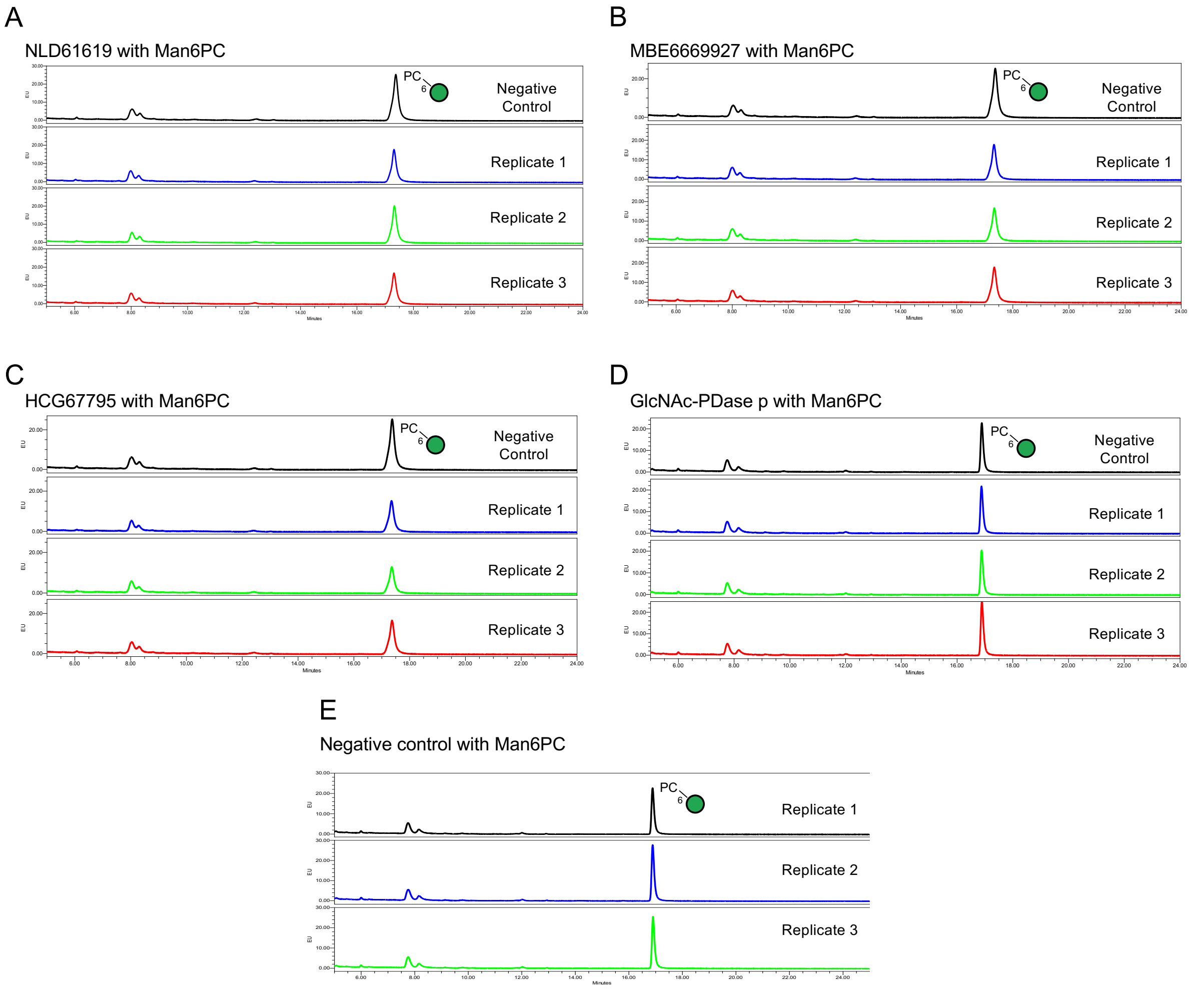


**Figure S9. GlcNAc-PDase and “Clostridia group” protein activity on Man6PC with UPLC-FLR analysis.**

IVTT-produced (A) NLD61619 (related protein 1), (B) MBE6669927 (related protein 2), (C) HCG67795 (related protein 3), (D) GlcNAc-PDase and (E) a negative control were incubated with the monosaccharide 6-O-phosphorylcholine-D-mannopyranose (Man-6-PC) in triplicate. Reactions were procainamide labeled and separated using UPLC-FLR.

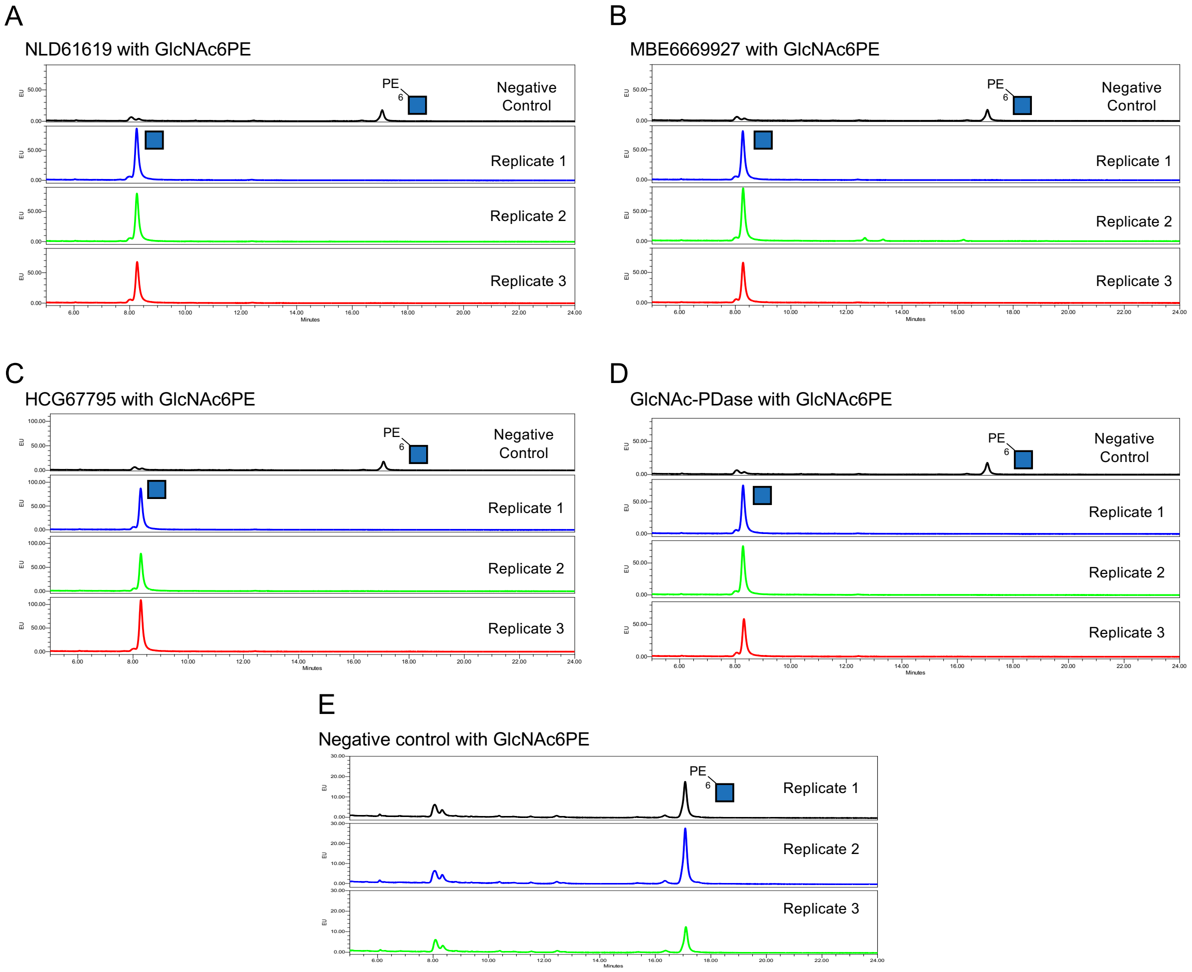


**Figure S10. GlcNAc-PDase and "Clostridia group" protein activity on GlcNAc6PE with UPLC-FLR analysis.**

IVTT-produced (A) NLD61619 (related protein 1), (B) MBE6669927 (related protein 2), (C) HCG67795 (related protein 3), (D) GlcNAc-PDase and (E) a negative control were incubated with the monosaccharide *N*-acetyl-D-glucosamine-6-phosphoethanolamine mannopyranose (GlcNAc-6-PE) in triplicate. Reactions were procainamide labeled and separated using UPLC-FLR.

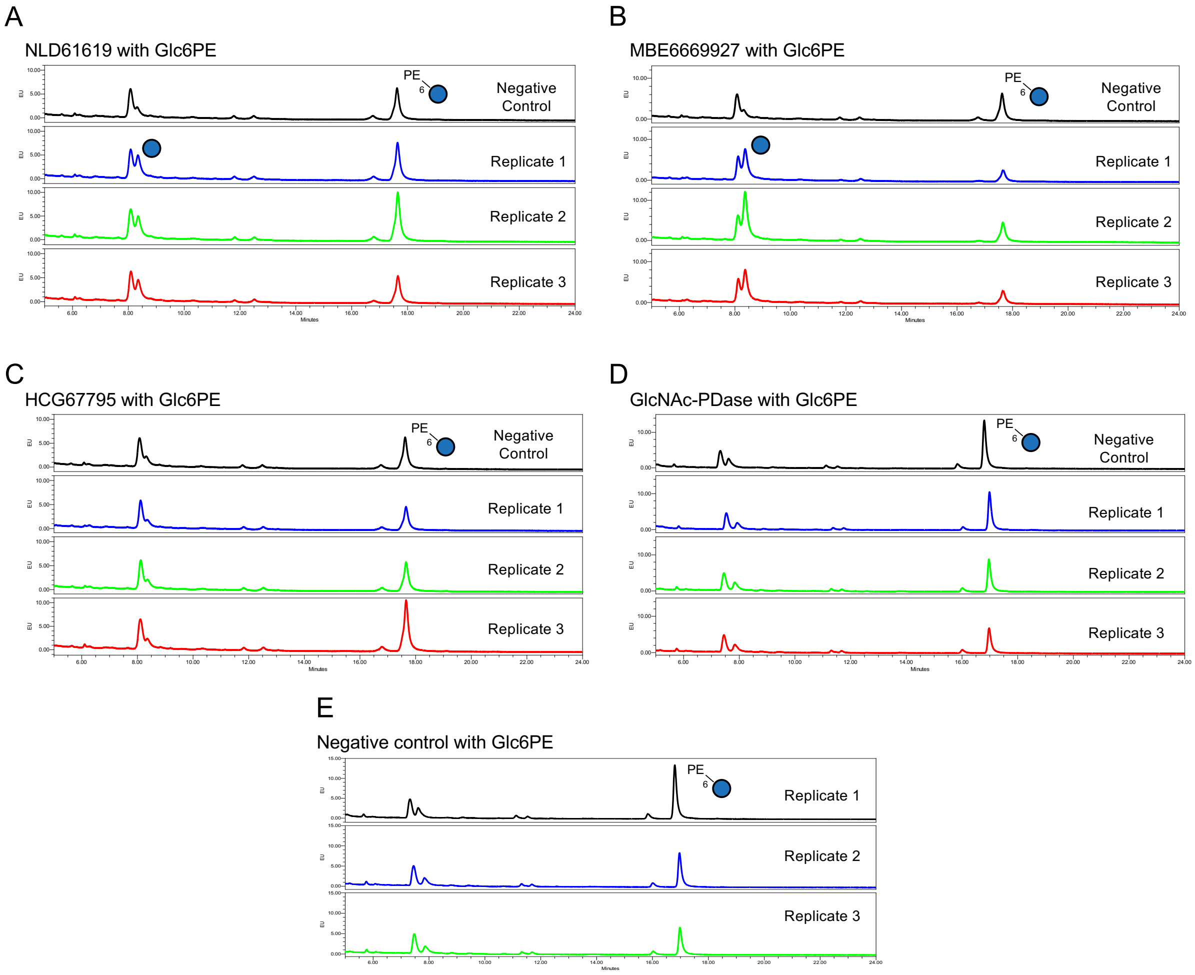


**Figure S11. GlcNAc-PDase and "Clostridia group" protein activity on Glc6PE with UPLC-FLR analysis.**

IVTT-produced (A) NLD61619 (related protein 1), (B) MBE6669927 (related protein 2), (C) HCG67795 (related protein 3), (D) GlcNAc-PDase and (E) a negative control were incubated with the monosaccharide 6-O-aminoethylphosphonato-D-glucopyranose (Glc-6-PE) in triplicate. Reactions were procainamide labeled and separated using UPLC-FLR.

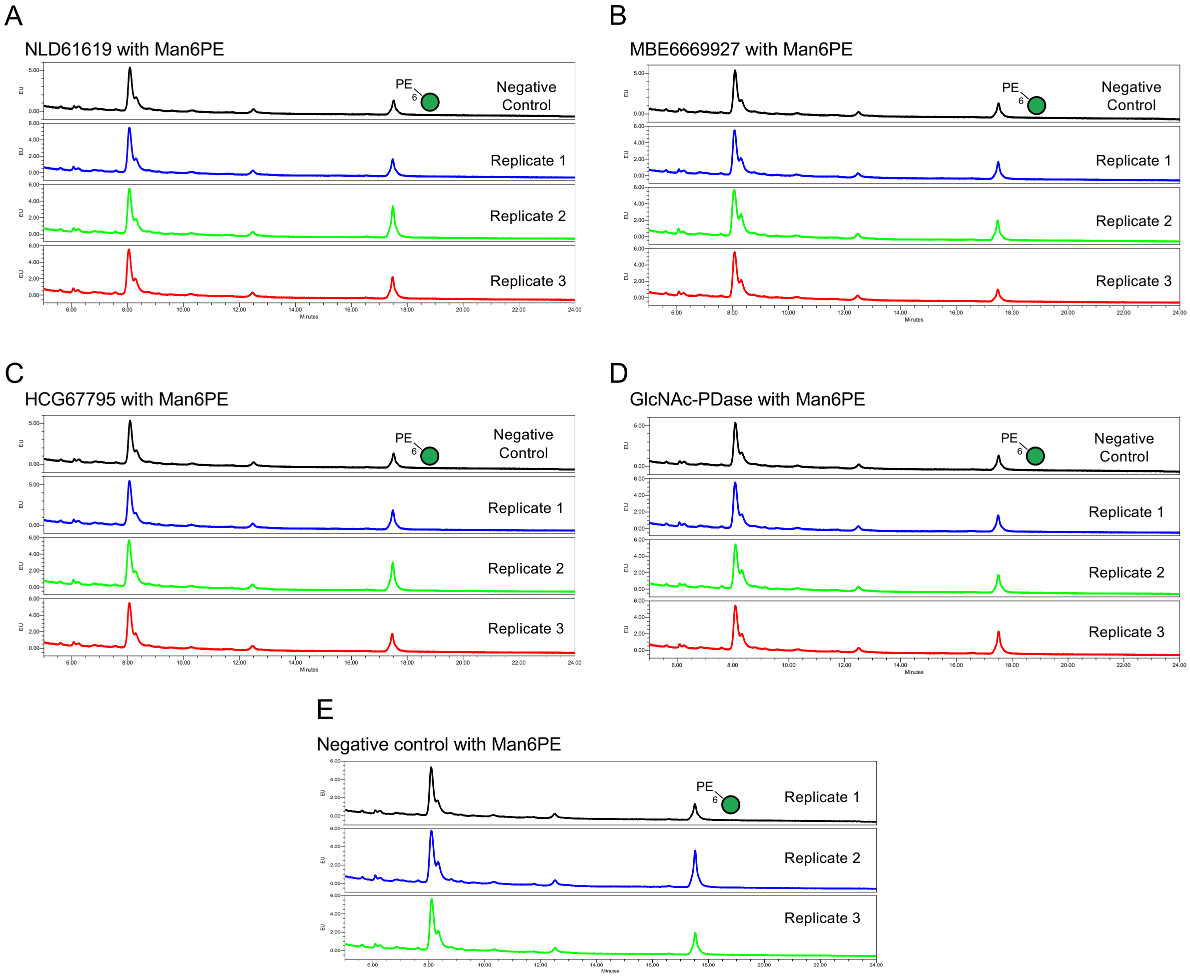


**Figure S12. GlcNAc-PDase and "Clostridia group" protein activity on Man6PE with UPLC-FLR analysis.**

IVTT-produced (A) NLD61619 (related protein 1), (B) MBE6669927 (related protein 2), (C) HCG67795 (related protein 3), (D) GlcNAc-PDase and (E) a negative control were incubated with the monosaccharide 6-O-aminoethylphosphonato-D-mannopyranose (Man-6-PE) in triplicate. Reactions were procainamide labeled and separated using UPLC-FLR.

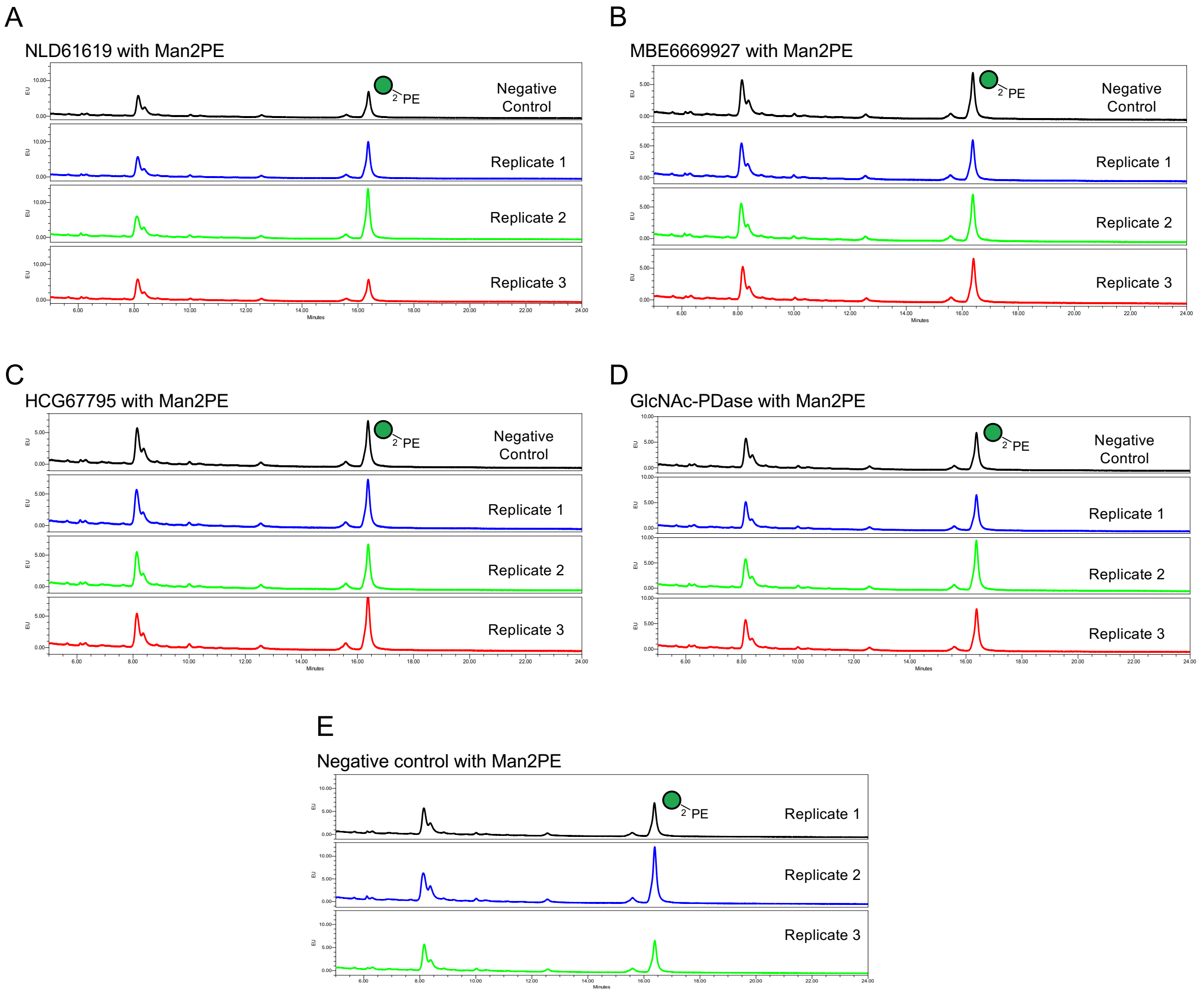


**Figure S13. GlcNAc-PDase and “Clostridia group” protein activity on Man2PE with UPLC-FLR analysis.**

IVTT-produced (A) NLD61619 (related protein 1), (B) MBE6669927 (related protein 2), (C) HCG67795 (related protein 3), (D) GlcNAc-PDase and (E) a negative control were incubated with the monosaccharide 2-O-aminoethylphosphonato-D-mannopyranose (Man-2-PE) in triplicate. Reactions were procainamide labeled and separated using UPLC-FLR.

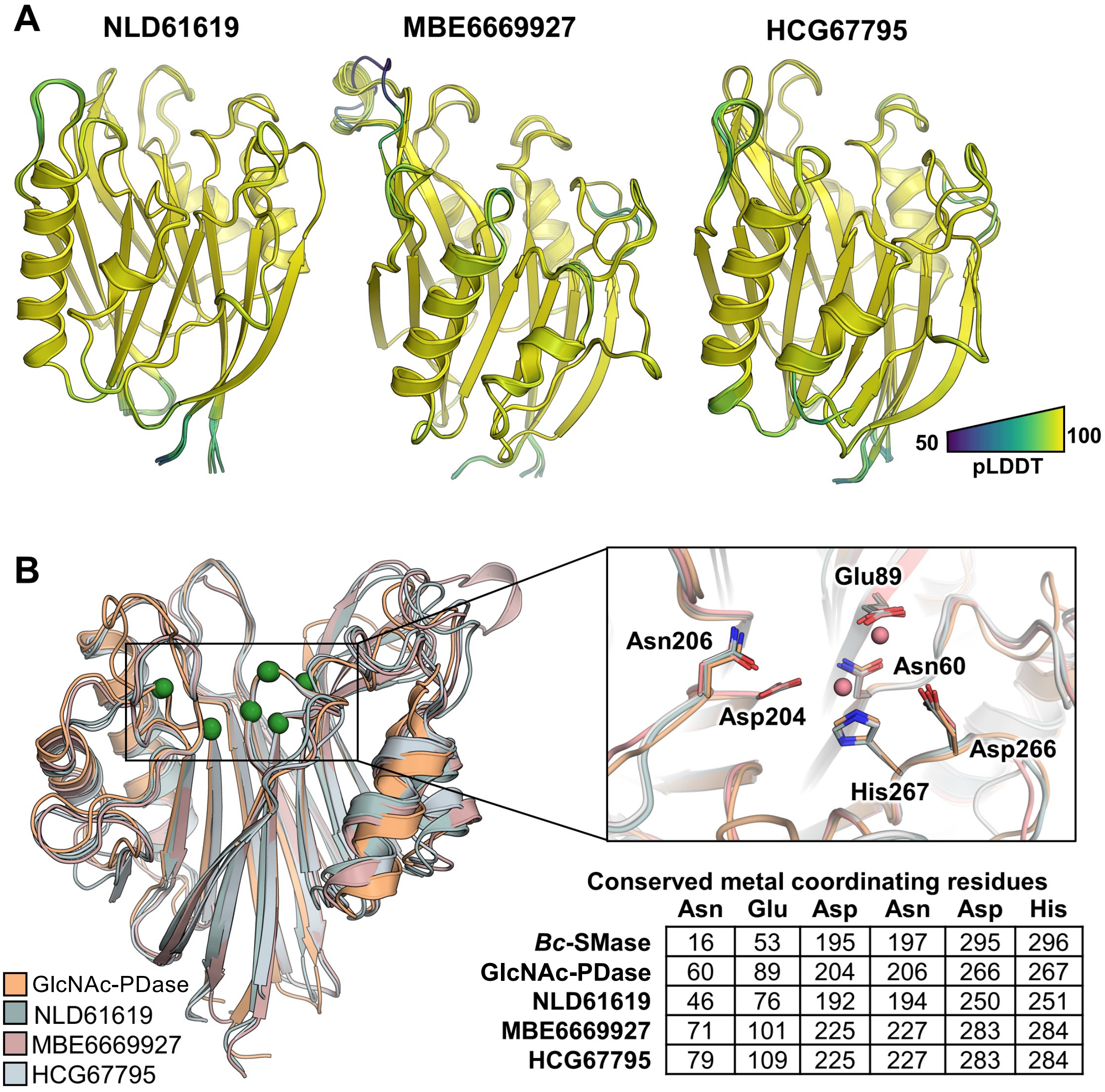


**Conserved metal coordinating residues**

| | Asn | Glu | Asp | Asn | Asp | His |
|---|---|---|---|---|---|---|
| ***Bc*-SMase** | 16 | 53 | 195 | 197 | 295 | 296 |
| **GlcNAc-PDase** | 60 | 89 | 204 | 206 | 266 | 267 |
| **NLD61619** | 46 | 76 | 192 | 194 | 250 | 251 |
| **MBE6669927** | 71 | 101 | 225 | 227 | 283 | 284 |
| **HCG67795** | 79 | 109 | 225 | 227 | 283 | 284 |

**Figure S14. Structural conservation analysis of GlcNAc-PDase with related proteins.**

A) Structural predictions of GlcNAc-PDase homologs NLD61619, MBE6669927, and HCG67795 in cartoon representation colored per residue by pLDDT score. The structured region of the top 5 ranks are shown superimposed to rank 1 by global alignment of C-alpha atoms.
B) Superposition of GlcNAc-PDase and related proteins demonstrate a consistent secondary structure folding pattern with apparent conserved metal coordinating residues at locations consistent with experimentally determined EEP enzyme *Bc*-SMase.

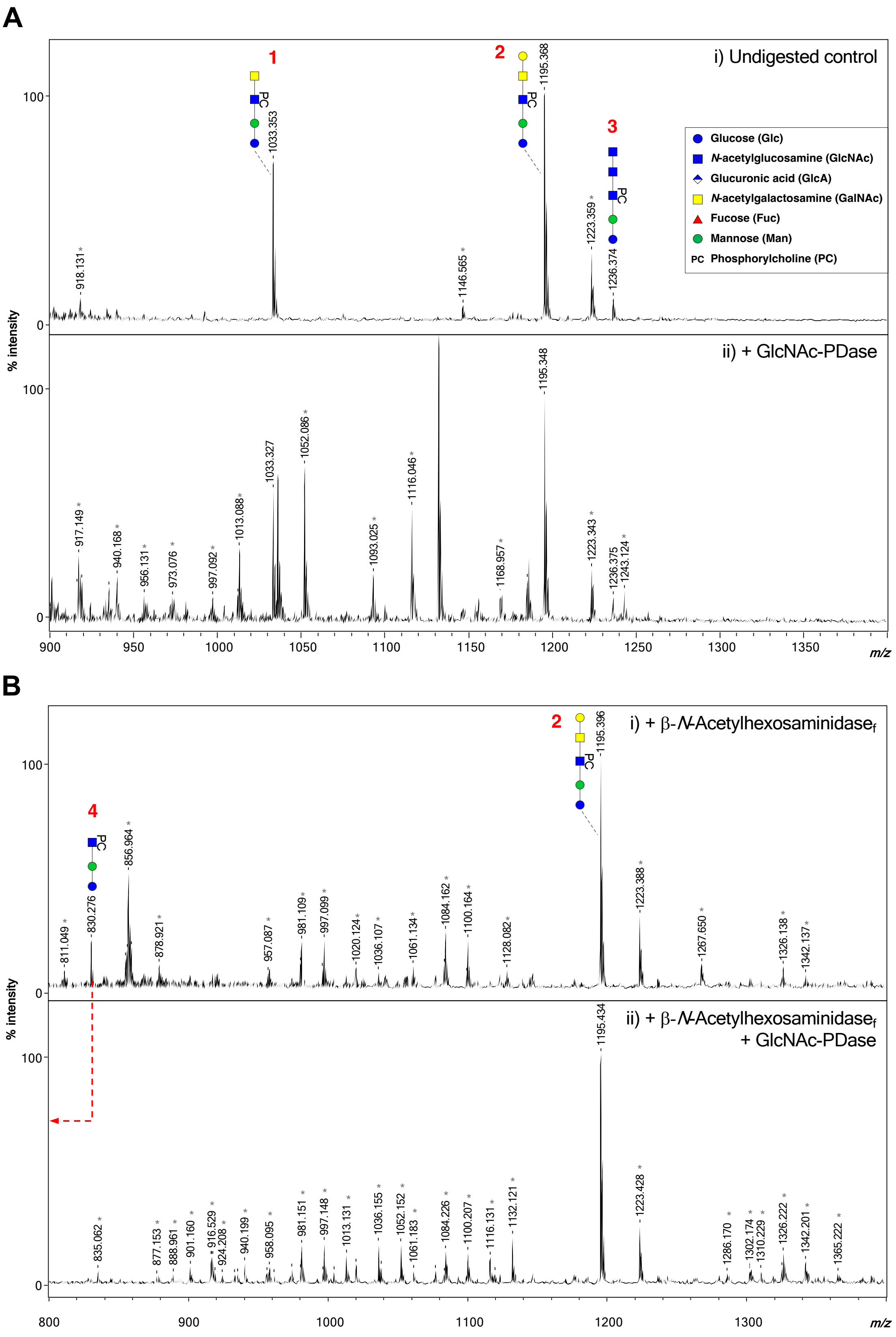
A
i) Undigested control
ii) + GlcNAc-PDase
B
i) + β-N-Acetylhexosaminidase_f
ii) + β-N-Acetylhexosaminidase_f + GlcNAc-PDase
% intensity
m/z
Glucose (Glc)
N-acetylglucosamine (GlcNAc)
Glucuronic acid (GlcA)
N-acetylgalactosamine (GalNAc)
Fucose (Fuc)
Mannose (Man)
PC Phosphorylcholine (PC)
1
2
3
4
PC
1033.353
1195.368
1223.359
1236.374
918.131
1146.565
1195.348
1052.086
1033.327
1116.046
1013.088
917.149
940.168
956.131
973.076
997.092
1093.025
1168.957
1223.343
1236.375
1243.124
1195.396
856.964
830.276
811.049
878.921
957.087
981.109
997.099
1020.124
1036.107
1061.134
1084.162
1100.164
1128.082
1223.388
1267.650
1326.138
1342.137
1195.434
1223.428
835.062
877.153
888.961
901.160
916.529
924.208
940.199
958.095
981.151
997.148
1013.131
1036.155
1052.152
1061.183
1084.226
1100.207
1116.131
1132.121
1286.170
1302.174
1310.229
1326.222
1342.201
1365.222

**Figure S15.** ***B. malayi*** **GSL glycans digestion with β-*N*-Acetylhexosominidase$_f$ and GlcNAc-PDase.**

Endoglucoceramidase (EGCase I; New England Biolabs)-released, AA-labeled and UPLC-purified *B. malayi* GSL glycans were subjected to digestion with GlcNAc-PDase (A.ii), β-*N*-Acetylhexosominidase$_f$ (B.i) or a combination of both enzymes (B.ii). Resulting digestion product is highlighted using a red dashed arrow. MALDI-TOF-MS spectra monoisotopic masses are indicated for ions with signal-to-noise ratio above 5. Non-glycan peaks are shown using grey stars and glycan structures are represented using the CFG nomenclature. See Symbol key inset. MALDI-TOF-MS raw data can be found in Table S2.B in the separate Excel file named "Table S2".

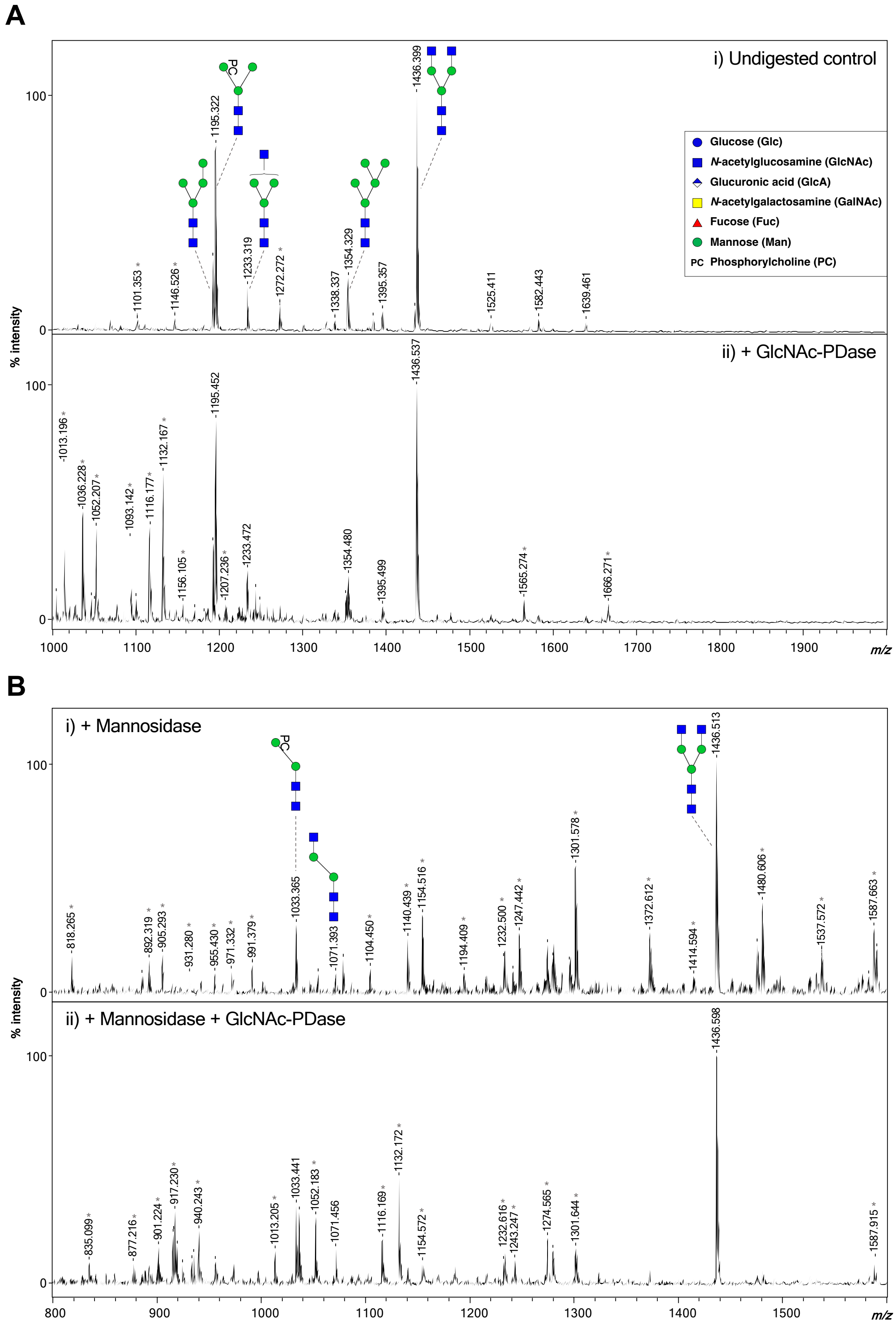
A
i) Undigested control
ii) + GlcNAc-PDase
Glucose (Glc)
N-acetylglucosamine (GlcNAc)
Glucuronic acid (GlcA)
N-acetylgalactosamine (GalNAc)
Fucose (Fuc)
Mannose (Man)
PC Phosphorylcholine (PC)
% intensity
m/z
B
i) + Mannosidase
ii) + Mannosidase + GlcNAc-PDase
% intensity
m/z

**Figure S16.** ***B. malayi*** **N-glycans digestion with α1-2,3,6 Mannosidase and GlcNAc-PDase.**

PNGase F-released, AA-labeled and UPLC-purified *B. malayi* N-glycans were subjected to digestion with GlcNAc-PDase (A.ii), α1-2,3,6 Mannosidase (B.i) or a combination of both enzymes (B.ii). MALDI-TOF-MS spectra monoisotopic masses are indicated for ions with signal-to-noise ratio above 5. Glycan structures are represented using the CFG nomenclature. See Symbol key inset. Non-glycan peaks are signaled using grey stars. MALDI-TOF-MS raw data can be found in Table S2.C in the separate Excel file named “Table S2”.

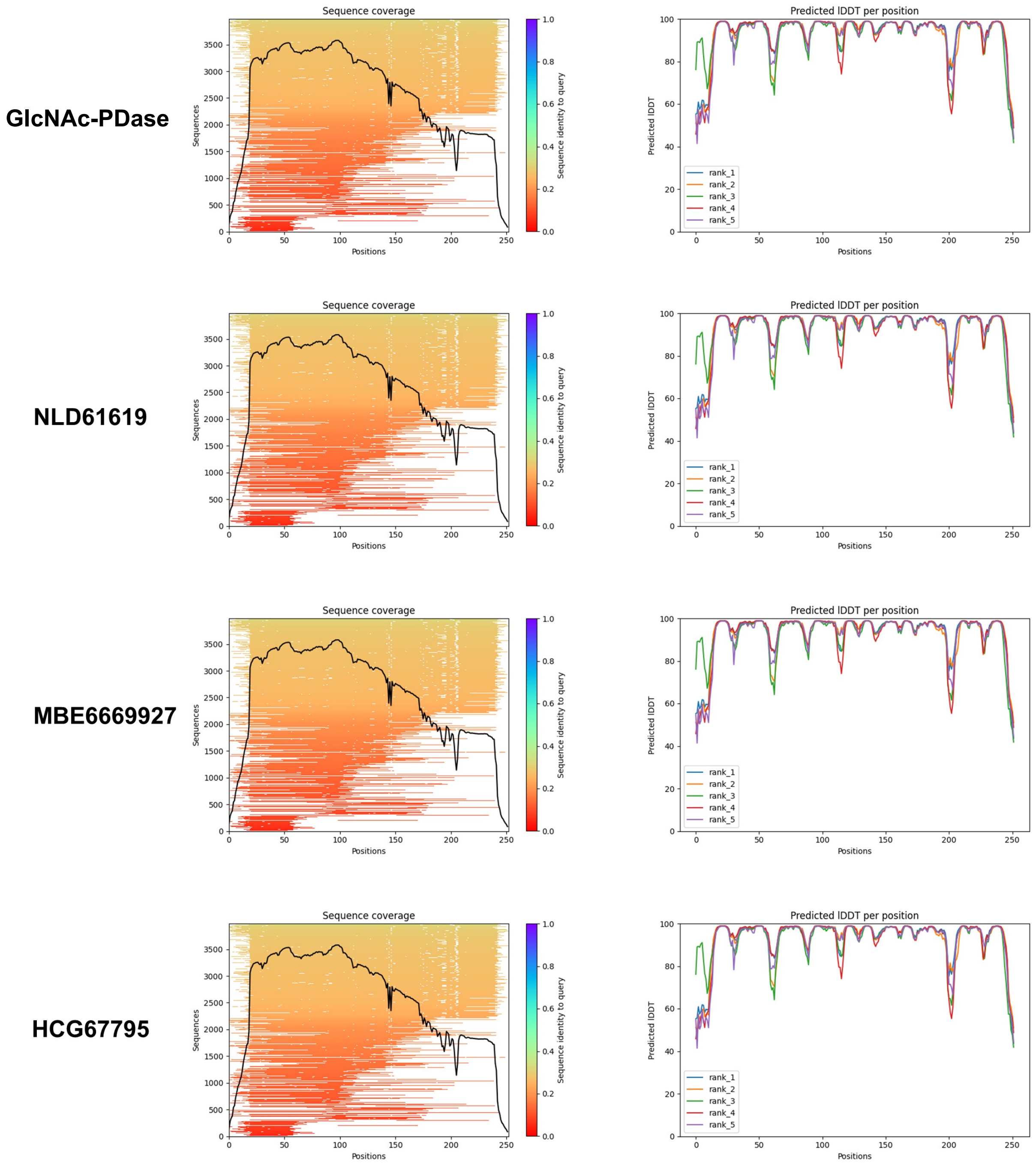


**Figure S17. MSA and pLDDT plots from ColabFold predictions.**
Multiple sequence alignment (MSA) coverage and Predicted Local Distance Difference Test (pLDDT) plots from ColabFold structural predictions for GlcNAc-PDase, NLD61619, MBE6669927, and HCG67795.

### Table S1. Primers used in this study.

| Primer | Sequence | Use |
|---|---|---|
| PURExpress_F1-ORF15_Fwd | GCGAATTAATACGACTCACTATAGGGCTTAAGTATAAGGAGGAAAAAATATGGAAGTAATACTCATTCGACACACT | *In vitro* expression of ORF1 from fosmid F1 |
| PURExpress_F1-ORF15_Rev | AAACCCCTCCGTTTAGAGAGGGGTTATGCTAGTTATTCCTTGTCGAGATTGATGCGGAC | *In vitro* expression of ORF1 from fosmid F1 |
| PURExpress_F2-ORF23_Fwd | GCGAATTAATACGACTCACTATAGGGCTTAAGTATAAGGAGGAAAAAATATGAAAGAGAACAAACCCTGCCTTTTG | *In vitro* expression of ORF23 from fosmid F2 |
| PURExpress_F2-ORF23_Rev | AAACCCCTCCGTTTAGAGAGGGGTTATGCTAGTTAATTTCCACATGATCTGTTTTCTAT | *In vitro* expression of ORF23 from fosmid F2 |
| PURExpress_F3-ORF6_Fwd | GCGAATTAATACGACTCACTATAGGGCTTAAGTATAAGGAGGAAAAAATATGGCATTTCTCGAAAGGAGCCCGGCT | *In vitro* expression of ORF6 from fosmid F3 |
| PURExpress_F3-ORF6_Rev | AAACCCCTCCGTTTAGAGAGGGGTTATGCTAGTTAAGCCTCCATCAGCACTTTCCCGTC | *In vitro* expression of ORF6 from fosmid F3 |
| PURExpress_F4-ORF27_Fwd | GCGAATTAATACGACTCACTATAGGGCTTAAGTATAAGGAGGAAAAAATATGGATATGTTGCAGCAAATTGGGAGC | *In vitro* expression of ORF27 from fosmid F4 |
| PURExpress_F4-ORF27_Rev | AAACCCCTCCGTTTAGAGAGGGGTTATGCTAGTTAAGCTTCATCCCGCTGCACCTCCGG | *In vitro* expression of ORF27 from fosmid F4 |
| PURExpress_F5-ORF14_Fwd | GCGAATTAATACGACTCACTATAGGGCTTAAGTATAAGGAGGAAAAAATATGAGCAAGGAATCCGTAACCGTCGCC | *In vitro* expression of ORF14 from fosmid F5 |
| PURExpress_F5-ORF14_Rev | AAACCCCTCCGTTTAGAGAGGGGTTATGCTAGTTATTGAGCCAGCAACCGGCGACCATA | *In vitro* expression of ORF14 from fosmid F5 |
| PURExpress_F5-ORF23_Fwd | GCGAATTAATACGACTCACTATAGGGCTTAAGTATAAGGAGGAAAAAATATGGCAGTGGTGTTGGCCGGCGTGGCA | *In vitro* expression of ORF23 from fosmid F5 |
| PURExpress_F5-ORF23_Rev | AAACCCCTCCGTTTAGAGAGGGGTTATGCTAGTTAGGCCAGTCCCATCTTGATTTTCAA | *In vitro* expression of ORF23 from fosmid F5 |
| PURExpress_F6-ORF3_Fwd | GCGAATTAATACGACTCACTATAGGGCTTAAGTATAAGGAGGAAAAAATATGAACGATAATAAAAACAGTATGAAA | *In vitro* expression of ORF3 from fosmid F6 |
| PURExpress_F6-ORF3_Rev | AAACCCCTCCGTTTAGAGAGGGGTTATGCTAGTTATAGCGCTCCCCTACGCAATATGTC | *In vitro* expression of ORF3 from fosmid F6 |

| | | |
|---|---|---|
| PURExpress_F7-ORF3_Fwd | GCGAATTAATACGACTCACTATAGGGCTTAAGTATAAGGAGGAAAAAATATGTCTGCAGCAAAAAGCAGTGACCTG | *In vitro* expression of ORF3 from fosmid F7 |
| PURExpress_F7-ORF3_Rev | AAACCCCTCCGTTTAGAGAGGGGTTATGCTAGTTATAAATATTTCACGCCCCAGACACT | |
| PURExpress_NLD61619_Fwd | GCGAATTAATACGACTCACTATAGGGCTTAAGTATAAGGAGGAAAAAATATGAAAGATGAGACAAAGTCAGAAATG | *In vitro* expression of NLD61619 |
| PURExpress_NLD61619_Rev | AAACCCCTCCGTTTAGAGAGGGGTTATGCTAGTTATTCAGACAATTTGGTTTGCGCATA | |
| PURExpress_MBE6669927_Fwd | GCGAATTAATACGACTCACTATAGGGCTTAAGTATAAGGAGGAAAAAATATGATTGATGAAGGAAATATATCAGAT | *In vitro* expression of MBE6669927 |
| PURExpress_MBE6669927_Rev | AAACCCCTCCGTTTAGAGAGGGGTTATGCTAGTTACTTAGAGCGGTATTTCAGGGTCGC | |
| PURExpress_HCG67795_Fwd | GCGAATTAATACGACTCACTATAGGGCTTAAGTATAAGGAGGAAAAAATATGTTACTATCAGCTTGTTCTGGAAGT | *In vitro* expression of HCG67795 |
| PURExpress_HCG67795_Rev | AAACCCCTCCGTTTAGAGAGGGGTTATGCTAGTTAGTCGCCGATGGACAGCGCCAACTC | |
| PURExpress_F4-ORF27mutant_N60A_E89A_N206A_D266A_H267A_Fwd | GCGAATTAATACGACTCACTATAGGGCTTAAGTATAAGGAGGAAAAAATATGGATATGCTACAACAGATAGGATCA | *In vitro* expression of ORF27 mutants from fosmid F4. Forward primer used to generate individual mutants for N60A, E89A, N206A, D266A, and H267A. |
| PURExpress_F4-ORF27mutant_N60A_Rev | AAACCCCTCCGTTTAGAGAGGGGTTATGCT AGTTACGCTTCATCACGCTGCACCTCCGG | *In vitro* expression of ORF27 mutant N60A from fosmid F4 |
| PURExpress_F4-ORF27mutant_E89A-Rev | AAACCCCTCCGTTTAGAGAGGGGTTATGCT AGTTATGCTTCATCACGCTGAACTTCCGG | *In vitro* expression of ORF27 mutant E89A from fosmid F4 |
| PURExpress_F4-ORF27mutant_D204A_Fwd | GCGAATTAATACGACTCACTATAGGGCTTAAGTATAAGGAGGAAAAAATATGGATATGCTACAACAGATAGGAAGT | *In vitro* expression of ORF27 mutant D204A from fosmid F4 |
| PURExpress_F4-ORF27mutant_D204A_Rev | AAACCCCTCCGTTTAGAGAGGGGTTATGCTAGTTATGCCTCATCACGTTGGACTTCCGG | |
| PURExpress_F4-ORF27mutant_N206A_Rev | AAACCCCTCCGTTTAGAGAGGGGTTATGCTAGTTAAGCCTCATCACGCTGCACTTCCGG | *In vitro* expression of ORF27 mutant N206A from fosmid F4 |
| PURExpress_F4-ORF27mutant_D266A_Rev | AAACCCCTCCGTTTAGAGAGGGGTTATGCTAGTTATGCTTCATCACGCTGAACTTCCGG | *In vitro* expression of ORF27 mutant D266A from fosmid F4 |

| | | |
|---|---|---|
| PURExpress-F4-ORF27mutant-H267A-R | AAACCCCTCCGTTTAGAGAGGGGTTATGCTAGTTACGCCTCATCACGCTGCACCTCCGG | *In vitro* expression of ORF27 mutant H267A from fosmid F4 |
| pJS119k_ORF27_Fwd | GAATTCAGCTTGGCTGTTTTG | Linearization of pJS119k vector for HiFi cloning of ORF27 |
| pJS119k_ORF27_Rev | ATGTTAACCTCCTAAGCTTAATTC | |
| F4-ORF27_Fwd | TTAAGCTTAGGAGGTTAACATATGGATATGTTGCAGCAAATTGGGAG | Amplifying ORF27 from fosmid F4 with C-terminal 8His for subcloning into pJS119k to generate pJS119k-GlcNAc-PDase-8His |
| F4-ORF27-8His(Cterm)_Rev | CAAAACAGCCAAGCTGAATTCTCAGTGGTGGTGGTGGTGGTGATGATGAGCTTCATCCCGCTGCACCTC | |

**Table S2. MALDI-TOF-MS raw data.**

This table can be found in the separate Excel file named “Table S2”. This file contains raw data for Supporting Information Figures S15 and S16, and Figure 6 of the main text.
